# AutoLEI: An XDS-based Graphical User Interface for Automated Real-time and Offline Batch 3D ED/MicroED Data Processing

**DOI:** 10.1101/2025.04.12.648515

**Authors:** Lei Wang, Yinlin Chen, Emma Scaletti Hutchinson, Pål Stenmark, Gerhard Hofer, Hongyi Xu, Xiaodong Zou

**Affiliations:** Department of Chemistry, Stockholm University, 106 91 Stockholm, Sweden; Department of Biochemistry and Biophysics, Stockholm University, 106 91 Stockholm, Sweden; Research School of Chemistry, the Australian National University, Acton, Australia

**Keywords:** 3D ED, MicroED, Real-time data processing, Offline batch data processing, Data analysis, Beam sensitive material

## Abstract

Three-dimensional electron diffraction (3D ED), also known as microcrystal electron diffraction (MicroED), is an emerging method for determining structures of submicron-sized crystals. With the development of rapid and convenient data collection protocols, acquiring dozens of datasets in a single MicroED session has become routine. A fast and automated workflow for processing, scaling and merging a large number of MicroED datasets can significantly accelerate the structure determination process. Herein, we present an XDS-based graphical user interface for <u>auto</u>mated rea<u>l</u>-tim<u>e</u> and offl<u>i</u>ne batch 3D ED/MicroED data processing (AutoLEI). We illustrate the functionality of the GUI through four examples, demonstrating both offline and real-time data processing capabilities. These examples include small organic molecules, metal-organic frameworks (MOFs), and proteins, showcasing the versatility and efficiency of the GUI in various applications.

**Synopsis:** A graphical user interface for real-time and offline 3D ED/MicroED data processing by XDS was developed. The GUI aims to improve efficiency, minimize redundant data processing work, and provide users with real-time feedback during data collection.

## 1. Introduction

Three-dimensional electron diffraction (3D ED), also known as micro-crystal electron diffraction (MicroED), has been developed into a robust method for crystal structure determination over the last few decades (Kolb *et al*., 2007; Zhang *et al*., 2010; Gemmi *et al*., 2019). The method enables rapid and accurate structure determination using commercially available transmission electron microscopes (TEM). A wide range of structures, including zeolite (Guo *et al*., 2015; Cho *et al*., 2023; Lu *et al*., 2024), metal-organic framework (MOF) (Huang, Willhammar *et al*., 2021; Huang, Grape *et al*., 2021), pharmaceuticals (Van Genderen *et al*., 2016; Lightowler *et al*., 2022), and more recently, macromolecules (Shi *et al*., 2013; Xu *et al*., 2019; Martynowycz *et al*., 2022), have been successfully solved with 3D ED/MicroED.

Benefiting from the strong interaction between electrons and matter, MicroED data can be collected from crystals that are too small for single-crystal X-ray diffraction. An ideal TEM grid typically contains hundreds of diffracting crystals or crystal fragments. With the latest MicroED protocol, it is possible to determine structures of high symmetry or stable samples through a single dataset within a few hours. However, due to the inherent limitations of the goniometer, data merging for crystals of low-symmetry is usually required to achieve high data completeness. For highly beam-sensitive material, a small-wedge data collection and merging strategy can also be employed to increase the signal-to-noise ratio and data completeness. Furthermore, selectively merging MicroED datasets can improve data quality by increasing redundancy (Xu *et al*., 2018; Samperisi *et al*., 2021; Bengtsson *et al*., 2022). More recently, high-throughput data collection has been achieved using software such as *Instamatic* (Cichocka et al., 2018), *Automated EM Data Acquisition with SerialEM* (De La Cruz *et al*., 2019), *Leginon* (Cheng et al., 2021), *EPUD* (Thermo Fisher Scientific), and *Latitude D* (GATAN) or dedicated electron diffractometer namely, *Synergy-ED* (Rigaku) and *ELDICO ED-1* (ELDICO Scientific). It is feasible to perform qualitative phase analysis by collecting and processing abundant datasets (Luo *et al*., 2023; Unge *et al*., 2024; Lightowler *et al*., 2024). In all the above cases, an efficient, automated, and even real-time data processing solution is highly desirable to enhance the efficiency of structure determination and phase analysis using MicroED.

A large selection of well-established software such as X-ray detector software (XDS) (Kabsch, 2010), PETS2 (Palatinus *et al*., 2019), CrysAlis^Pro^-ED (Truong *et al*., 2023), xia2.multiplex (Gildea *et al*., 2022) and DIALS (Winter *et al*., 2018, 2022), are available for processing MicroED data. Data processing can be performed via either command-line instructions or graphical user interfaces (GUI) (Brehm *et al*., 2023), offering flexibility to users. However, format conversion, sorting, processing, analyzing and merging a large amount of MicroED datasets can be time-consuming and laborious. Automated batch processing and real-time processing capabilities are still underdeveloped for MicroED. In our previous work, Edtools (Wang *et al*., 2019) was developed to process batch datasets in offline mode but only works in the command line. A processing pipeline Scipion-ED (Bengtsson *et al*., 2022) embedded in Scipion was also attempted. However, the complex operation and specialized platform requirement make them difficult for beginners to use.

In this work, we utilized XDS as the data processing engine and developed a graphical user interface for <u>Auto</u>mated Real-tim<u>e</u> and Offline Batch 3D ED/MicroED Data Processing (AutoLEI). Based on the fast data processing performance of XDS, AutoLEI enabled both offline and real-time batch MicroED data processing. This Python-based GUI is straightforward to install and provides users with the options and parameters that are essential for working with electron diffraction data. Meanwhile, data processing statistics and quality indicators are also optimized for MicroED. We demonstrate how to use AutoLEI to process data collected from different types of crystals, including an organic molecule, a metal-organic framework and two proteins, on different microscopes and detectors. An example of real-time data processing using AutoLEI is also presented. Furthermore, a detailed tutorial with training datasets is provided.

## 2. AutoLEI implementation

### 2.1. AutoLEI installation and data format supporting

AutoLEI is an open-access GUI written in Python. It can be downloaded from PyPI, Gitlab or Zenodo. In version 1.0.0 the native image format is SMV for XDS data processing. Automated image format conversion supports TIFF images collected on CheeTah (Amsterdam Scientific Instruments), OneView (GATAN), Timepix hybrid pixel detectors (Amsterdam Scientific Instruments) and MRC images collected by EPUD (Thermo Fisher Scientific). Real-time data processing is designed for raw MRC image data collected using EPUD (Thermo Fisher Scientific) as well as data collected using *Instamatic*. Future updates are expected for improved compatibility and increased support, while users can also add new functionality through the programming interface.

### 2.2. AutoLEI data processing workflow

The primary aim of AutoLEI is to process and analyze a batch of MicroED datasets using XDS offline, or in real-time. Figure 1 shows the workflow of AutoLEI. Data processing goes through the steps of ***XDSRunner, CellCorr, XDSRefine, MergeData*** and ***Cluster&Output***. These steps are implemented as tabs in the AutoLEI interface as shown in Supporting information (SI) Figures S1–S8. By following this workflow, final HKL files are obtained for downstream structural determination. Videos 1–3 (SI) showcase the offline data processing of three examples: tyrosine, MOF SU-100 and the protein *H. sapiens* MutT homolog 1 (hMTH1) while Video 4 (SI) shows the real-time data processing of triclinic lysozyme.

**Figure 1.**
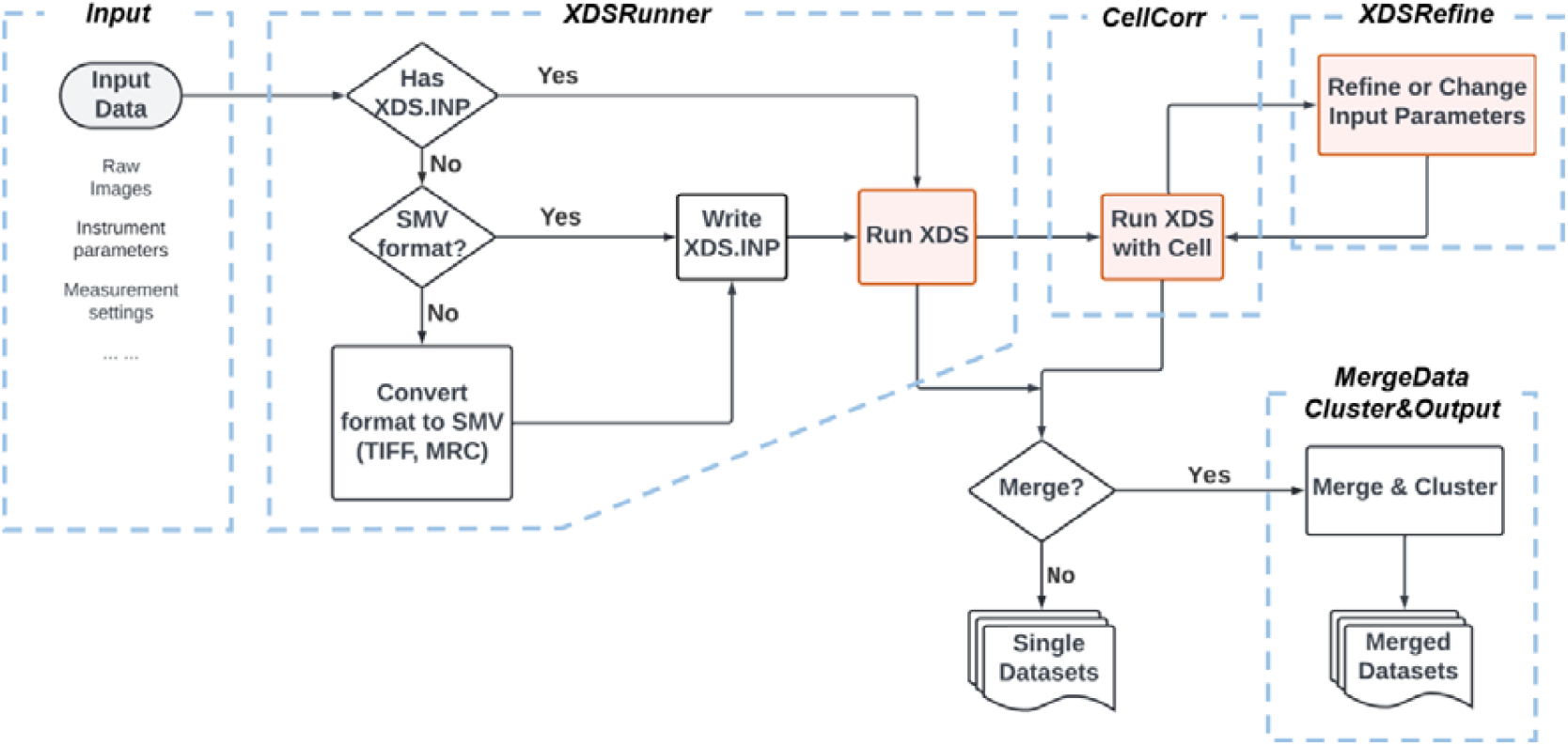
An overview of the AutoLEI workflow

#### 2.2.1. Input

***Input*** allows users to set up the work directory, instrument parameters, and measurement settings (Figure S1). The instrument parameters can be loaded from a predefined instrument model or existing input files. Alternatively, they can be given manually when datasets collected from different instruments and detectors are used. ***Input*** creates the “Input_parameters.txt” file, which will be used for generating input files “XDS.INP” for both real-time and offline MicroED data processing.

#### 2.2.2. XDSrunner

***XDSRunner*** generates the corresponding “XDS.INP” files for each dataset (Figure S2). It then launches XDS to process these datasets. No prior knowledge of the sample is required. AutoLEI processes all datasets sequentially under the work directory and parses statistics from XDS outputs. After data processing, AutoLEI will generate a summary table named “xdsrunner.xlsx”. The table contains key parameters and quality indicators such as space group (SG), unit cell parameters (Unit cell), cell volume (Vol.), indexing rate (Index%), I/Sigma(I)^asymptotic^ (ISa), redundancy-independent merging R-factor (*R*_meas_), correlation between reflection intensities of random half datasets (*CC*_1/2_), completeness and estimated resolution (Reso.) of each dataset for inspection. SG, unit cell and ISa are read from XDS outputs directly while others are calculated by AutoLEI based on XDS HKL files. The high-resolution cut-off of each dataset is estimated by checking the *CC*_1/2_, *R*_int_ and signal-to-noise ratio of each resolution shell.

***XDSRunner*** also contains supporting functions as shown below:

- *Format converter*: Convert raw image files to SMV and write the metadata into SMV files.
- *Find beam center*: Find the beam center automatically with or without a beam stop.
- *Estimate symmetry & cell-cluster*: Cluster data based on unit cell parameters and estimate the Laue group. This ensures data can be processed in the correct Laue group and unit cell.
- Details of these functions can be found in Supporting Information (S1. Supporting function implementation in AutoLEI).

#### 2.2.3. CellCorr

***CellCorr*** allows users to run XDS with given unit cell parameters and space group (Figure S3). It allows re-indexing the original HKL using space group and unit cell parameters suggested by either “Estimate symmetry & cell-cluster” in ***XDSRunner*** or determined by other software packages such as REDp (Wan *et al*., 2013) and PETS2. This ensures that the datasets will be processed, scaled and merged in the correct Laue group and unit cell. ***CellCorr*** outputs a summary table named “xdsrunner2.xlsx” containing the same data quality indicators as “xdsrunner.xlsx”.

#### 2.2.4. XDSRefine

***XDSRefine*** provides the opportunity to update XDS keywords in the input files for single, selected, or all datasets if users are not satisfied with the outputs from ***CellCorr*** (Figure S4). The step is useful for problematic data that cannot be processed using default settings. In AutoLEI v1.0.0, ***XDSRefine*** also provides automatic refinement of the rotation axis, and removal of scale outliers caused by crystal shifts during data collection. The reconstructed reciprocal lattice of each dataset can also be visually inspected. ***XDSRefine*** updates “xdsrunner2.xlsx” and the table will then be used for downstream data scaling and merging.

#### 2.2.5. MergeData

***MergeData*** analyzes statistics in “xdsrunner2.xlsx” and make initial selections of datasets for merging (Figure S5). Users can then inspect key indicators listed in the table and apply a filter for scaling/merging. The default value will be provided as a reference. The information on filtered data is written in “xdspicker.xlsx”. ***MergeData*** will then initiate XSCALE and output scaled data as an HKL file.

#### 2.2.6. Cluster&Output

***Cluster&Output*** performs cluster analysis based on pair-wise correlation coefficients of reflection intensities in “XSCALE.LP” generated from the last step (Figure S6). A dendrogram will be produced for visual inspection. Users can set the distance threshold for further clustering. ***Cluster&Output*** will then perform scaling/merging of each cluster, output corresponding “HKL” files and data processing statistics (Figure S7). Webpage-based reports will also be generated for inspection of data reduction.

### 2.3. Real-time processing

Real-time MicroED data processing is a fully automated pipeline in AutoLEI (Figure S8). With default or customized settings, it provides crucial feedback to users for data quality analysis, evaluates data collection parameters, and generates result “HKL” files in real-time. The setup of the real-time processing function is found on the ***RealTime*** tab of AutoLEI, as shown in Figure 2. Real-time processing reads “Input_parameters.txt” from ***Input*** and continuously monitors the input folder where the data is being saved. Once a single dataset collection is completed, real-time processing will try to process data in either screening mode or merging mode:

a. *Screening mode*: When unit cell parameters and space group are not provided, the data is processed without performing data merging.
b. *Merging mode*: When unit cell parameters and space group are provided, the data will be processed in the specified cell and space group. If the data quality satisfies the filtering condition, it will be assigned as a valid dataset and merged with other valid datasets.

**Figure 2.**
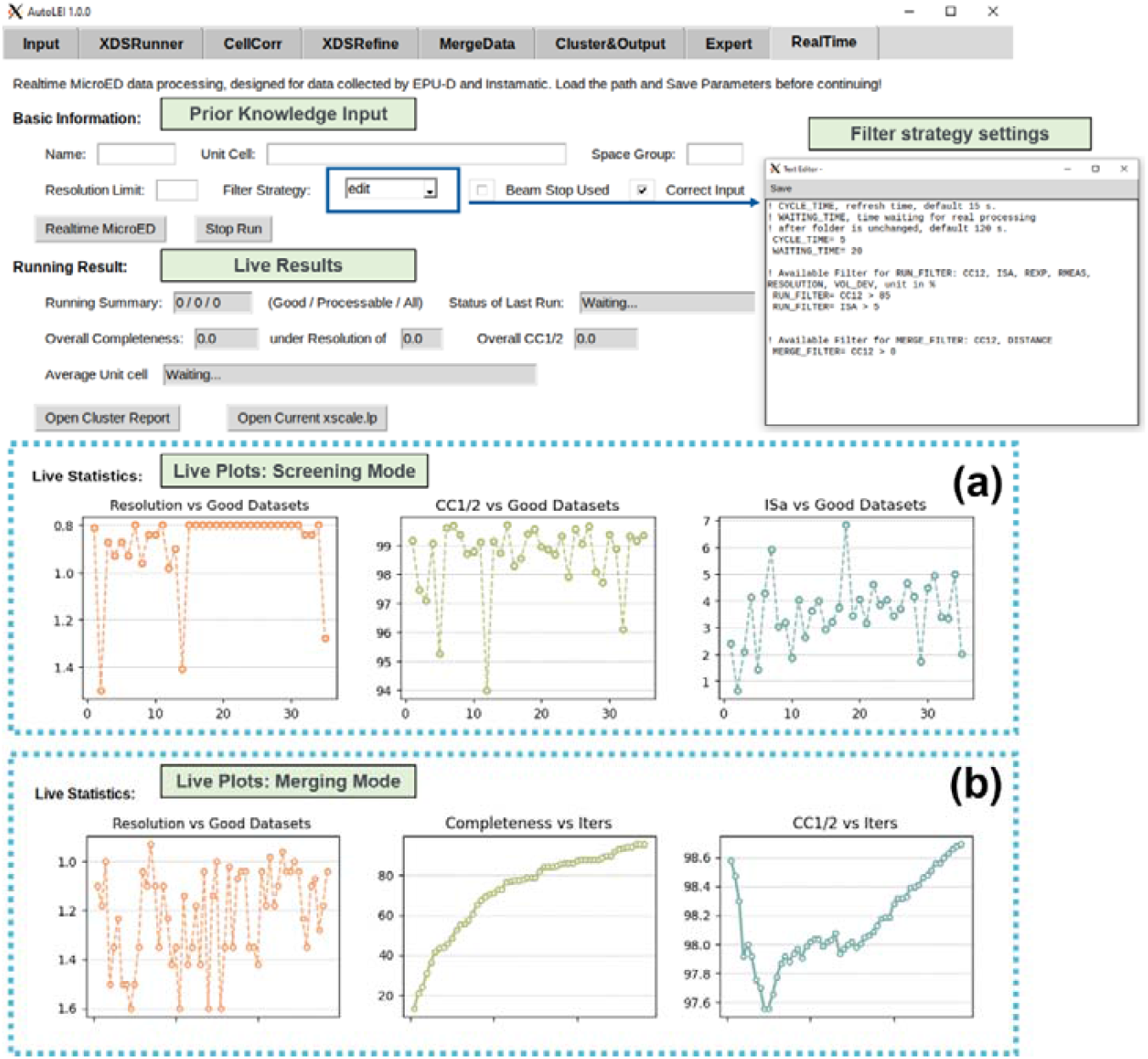
Real-time MicroED work page. (a) Screen mode. (b) Merge mode.

In the ***RealTime*** tab, estimated resolution, *CC*_*1/2*_ and ISa for individual data or merged data are plotted and updated live, as shown in Figure 2a and 2b, respectively. Real-time processing currently supports the data collected using EPUD and Instamatic.

## 3. Examples

Table 1 and Figure 3 summarize four examples of data processing using AutoLEI. MicroED data were collected on a variety of microscopes and detectors using different software. For sample preparation details, please refer to the Supporting Information (S2). Among them, tyrosine (Video 1), SU-100 (Video 2) and hMTH1 (Video 3) use offline data processing protocol, while lysozyme (Video 4) showcases the online data processing capability.

**Table 1.** Summary of data used for demonstrating AutoLEI processing.

| Sample | Microscope | Detector | Data collection software | Total datasets | Data collection time | Data processing time<sup>1</sup> |
| --- | --- | --- | --- | --- | --- | --- |
| Tyrosine<br>(P2<sub>1</sub>2<sub>1</sub>2<sub>1</sub>) | JEM-2100<br>LaB6 (JEOL) | Timepix hybrid<br>pixel detector (ASI) | *Instamatic* | 12 | ~ 1 h 20 min | ~ 5 min |
| SU-100 (C2/c) | JEM-2100<br>LaB6 (JEOL) | Timepix hybrid<br>pixel detector (ASI) | *Instamatic* | 16 | ~ 2 h 15 min | ~ 7 min |
| hMTH1<br>(P2<sub>1</sub>2<sub>1</sub>2<sub>1</sub>) | Krios G3i<br>(TFS) | Ceta-D (TFS) | EPUD (TFS) | 38 | ~ 2 h 14 min | ~ 12 min |
| Lysozyme (P1) | Krios G2<br>(TFS) | Ceta-D (TFS) | EPUD (TFS) | 72 | ~ 6 h | Real-time |

<sup>1</sup> Timing is from the start of data processing to obtaining usable “HKL” files (Videos 1–3)
JEOL: Japan Electron Optics Laboratory Co., LTD
ASI: Amsterdam Scientific Instruments
TFS: Thermo Fisher Scientific

**Figure 3.**
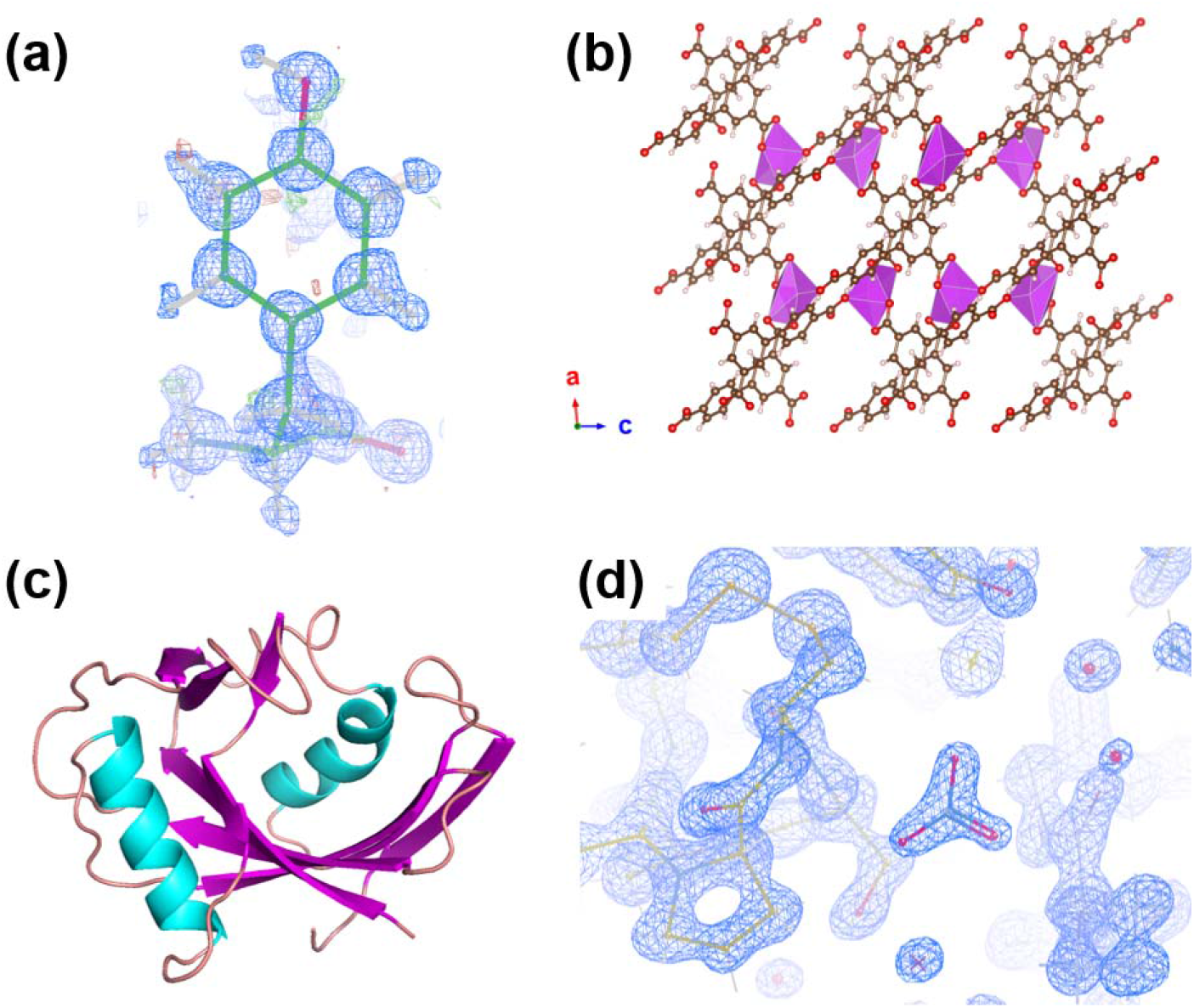
Summary of solved structures. (a) The structure of L-tyrosine using data from experiment_10. Electrostatic potential map (blue) and difference map (green and red) and tyrosine structure shown in COOT(Emsley *et al*., 2010). The electrostatic potential map (blue) was contoured at 1σ-level while the difference map (red and green) was countered at 3σ-level. (b) The structure framework of SU-100 viewed along the *b*-axis. (c) The structure of apo-hMTH1. (d) No-filled 2*F*o−*F*c electrostatic potential map (blue) with lysozyme model near a nitrate anion. The map was contoured at 1.5 r.m.s.d.

### 3.1. Single-data-based structure determination: rapid filtering of low-quality tyrosine data

Tyrosine is a naturally occurring amino acid and an essential protein building block. By collecting MicroED data over a large tilt range, structure determination from a single orthorhombic crystal is feasible. A total of 12 tyrosine datasets were collected and then processed with a default resolution range from 20 to 0.8 Å. Unit cell parameters, ISa, *CC*_1/2_, completeness and estimated resolutions were determined and extracted from XDS outputs. As shown in Figure 4a, most of the datasets displayed favorable data quality and the estimated resolution reached the pre-defined high-resolution limit (0.8 Å) in ***XDSRunner*** tab. This suggested that the resolution range should be extended in the following procedure. However, both experiment_3 and experiment_7 suffered from unacceptable low completeness because of the low tilting range during data collection, indicating that these datasets might not be suitable. Subsequently, the unit cell parameters and Laue group were extracted from the output of “Estimate symmetry & cell clustering” and then used for XDS processing in ***CellCorr. XDSrefine*** to refine the rotation axis, divergence and mosaicity, scale outliers and change resolution range to 30–0.5 Å, as shown in Figure 4b.

**Figure 4.**
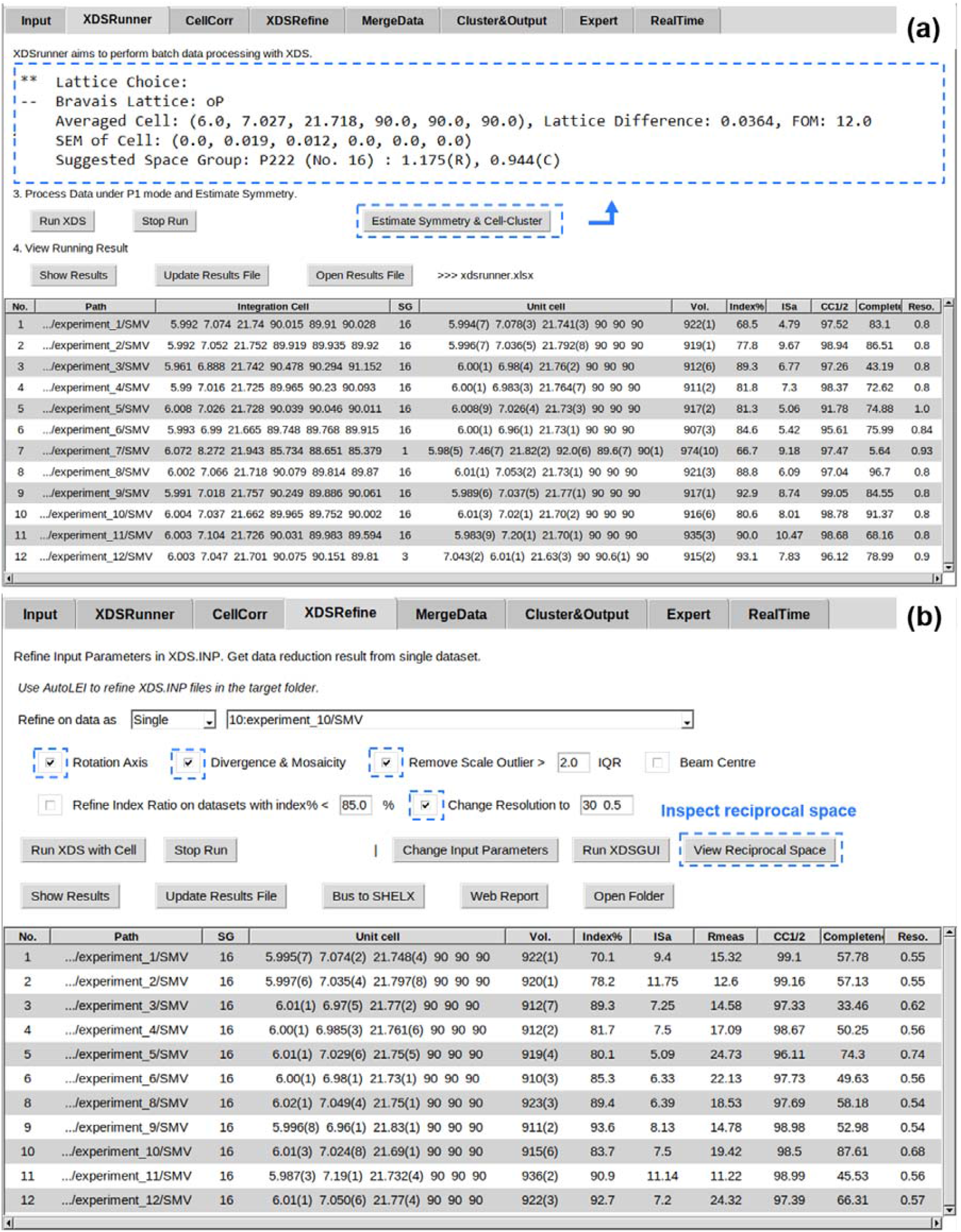
Datasets of 12 L-tyrosine crystals processed using AutoLEI. (a) Result table after running ***XDSrunner***. The suggested unit cell parameters and space group were extracted from the “Estimate Symmetry & Cell Cluster” function. (b) Result table after running ***XDSRefine***. “View Reciprocal Space” was designed to inspect the reciprocal space of a single dataset.

Different users may prioritize different key indicators when selecting the optimal datasets. Experientially, we recommend datasets with an ISa > 5, *CC*_1/2_ > 95%, and completeness > 80% as being promising for single-data-based structure determination. We here selected experiment_10 due to its good data quality: Isa, *CC*_1/2_, completeness and estimated resolution of this data were 7.51, 98.18%, 87.61% and 0.68 Å, respectively. AutoLEI also enables the systematic absence in reciprocal space to be visually inspected, as shown in Figure S9. It is noted that the 0*k*0 row was missing in most of datasets due to preferential crystal orientation. By using AutoLEI, it was possible to identify the missing 0*k*0 row in experiment_12 by checking the reciprocal space of all data rapidly. The space group could then be deduced as *P*2_1_2_1_2_1_. The time used for data processing and analysis was approximately 5 minutes in total. The integrated data was then used for structure solution and kinematic refinement. The number of unique reflections and parameters used for refinement was 2592 and 162, respectively. The final *R*_1_ converged to 11.90% (Table S1). All hydrogen atoms including the proton near the N atom could be found as shown in Figure 3a.

To further investigate the relationship between data quality and the structure information, we solved the structure using other datasets, as shown in Table S1. Except for experiment_3 and experiment_7, which had low completeness, all other datasets could be used for successful structure determination.

### 3.2. Interactive data merging of MOF SU-100

SU-100 is a monoclinic MOF material, and its flexible framework exhibits promising breathing behaviors (Grape *et al*., 2020). In this study, a total of 16 datasets were collected using Instamatic, and all data were processable after ***XDSRefine*** as shown in Figure 5a. Data with an ISa higher than 5 were then merged during the ***XDSRefine*** step after specifying the space group and unit cell parameters (Figure 5b). We further conducted cluster analysis based on the pair-wise correlation coefficients of reflection intensities. As shown in Figure 5c, a threshold of 0.5 was selected due to the significant distance between four clusters. The final “HKL” file was obtained in 7 minutes. The resolution cut-off was 0.62 Å. As shown in Figure 3b, all non-H atoms could be directly determined from the solution. The number of unique reflections and parameters used for refinement was 7193 and 200, respectively. The final *R*_1_ converged to 19.46% as shown in Table S2.

**Figure 5.**
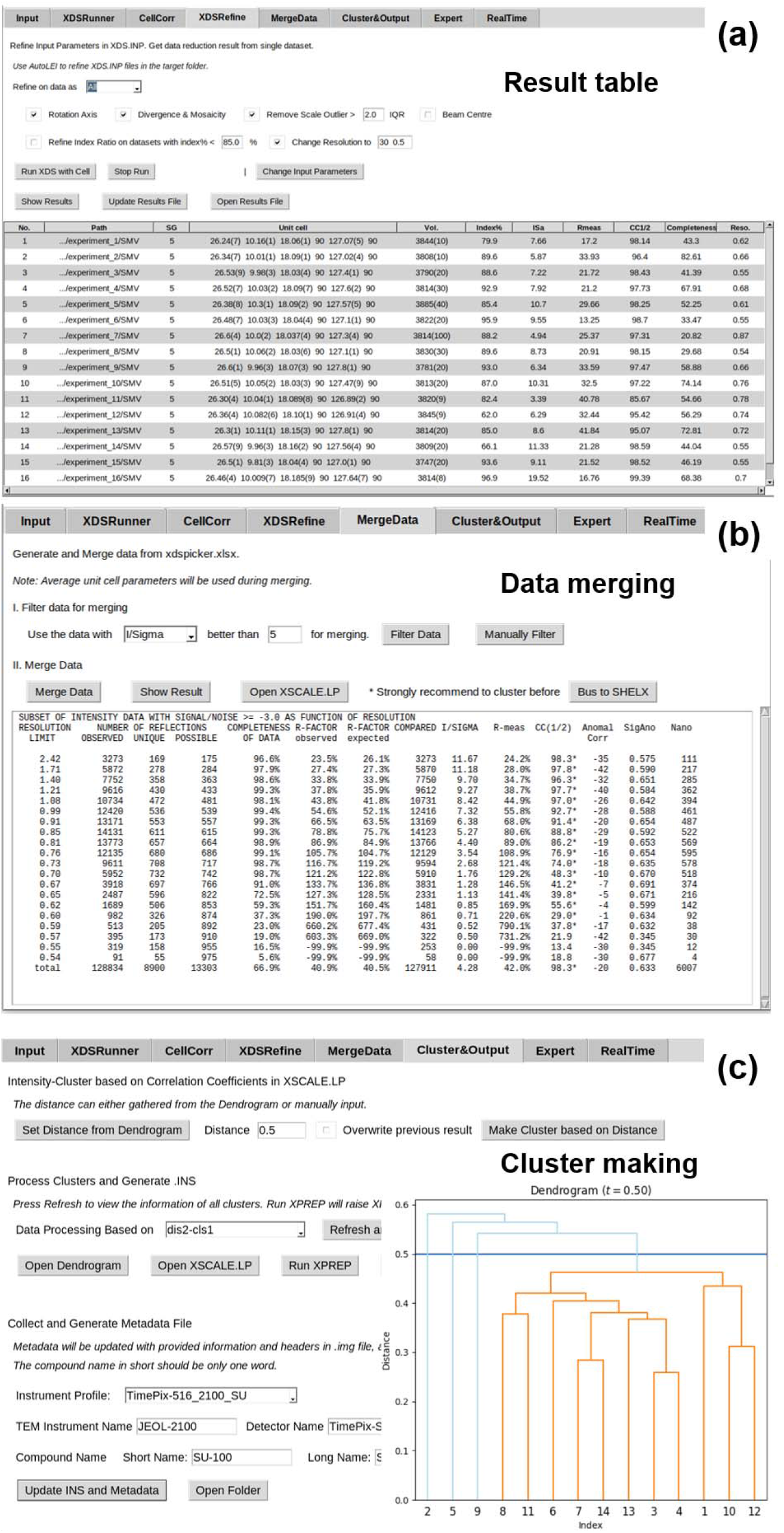
Data processing, analysis and merging of 16 SU-100 MOF datasets by AutoLEI. (a) ***XDSRefine*** work page showing the results table. (b) ***MergeData*** work page. The initial round of data merging was achieved after several clicks. (c) ***Cluster_Output*** work page. The clusters based on pair-wise reflection intensity correlations were made, and a final “HKL” file was prepared for structure determination.

### 3.3. Processing small wedge MicroED data of protein MutT homolog 1 (hMTH1)

Protein crystals are known for being extremely beam-sensitive during MicroED data collection (Shi *et al*., 2013). To address this issue, a small-wedge data collection strategy was adopted for MicroED data collection. However, this strategy produces a large number of datasets that are of low completeness individually. Thus, repetitive data processing, data scaling and merging are required for structure determination, which is an ideal application for AutoLEI.

hMTH1 plays an important role in sanitizing oxidized dNTPs from the free nucleotide pool, preventing their incorporation into DNA, which reduces genotoxicity (Yoshimura *et al*., 2003; Jemth *et al*., 2018). Here, we used hMTH1 as an example to demonstrate how small wedge MicroED data can be effortlessly processed and merged with AutoLEI. A total of 38 datasets were collected using EPUD and the average rotation range of each crystal was limited to 15-20 degrees to boost the signal-to-noise ratio while reducing beam damage. Figure 6a shows the dendrogram of the clustering analysis after the ***XDSRefine*** step. AutoLEI also enables a stepwise clustering analysis, which is useful when processing protein data. Different distance thresholds were tested in order to find the optimal distance threshold. Based on this analysis, the first four groups containing 27, 26, 23 and 18 datasets showed comparable completeness while the completeness of the last group (contained 13 datasets) dropped by more than 4%. Thus, Cluster 1 based on threshold 4 (dis4-cls1) was selected for higher *CC*_*1/2*_ and lower *R*_meas_. A web report was then generated and used for data reduction of this cluster. Figure 6b indicates the relationships between *CC*_1/2_, *R*_meas_ and completeness against resolution printed from the report of dis4-cls1. The estimated resolution was 2.76 Å and it took less than 12 minutes to obtain the final “HKL” file using AutoLEI. Molecular replacement and structure refinement were carried out using the Phenix package (Liebschner *et al*., 2019). As shown in Table S3, the final *R*_work_ and *R*_free_ converged to 21.07% and 26.82%, respectively.

**Figure 6.**
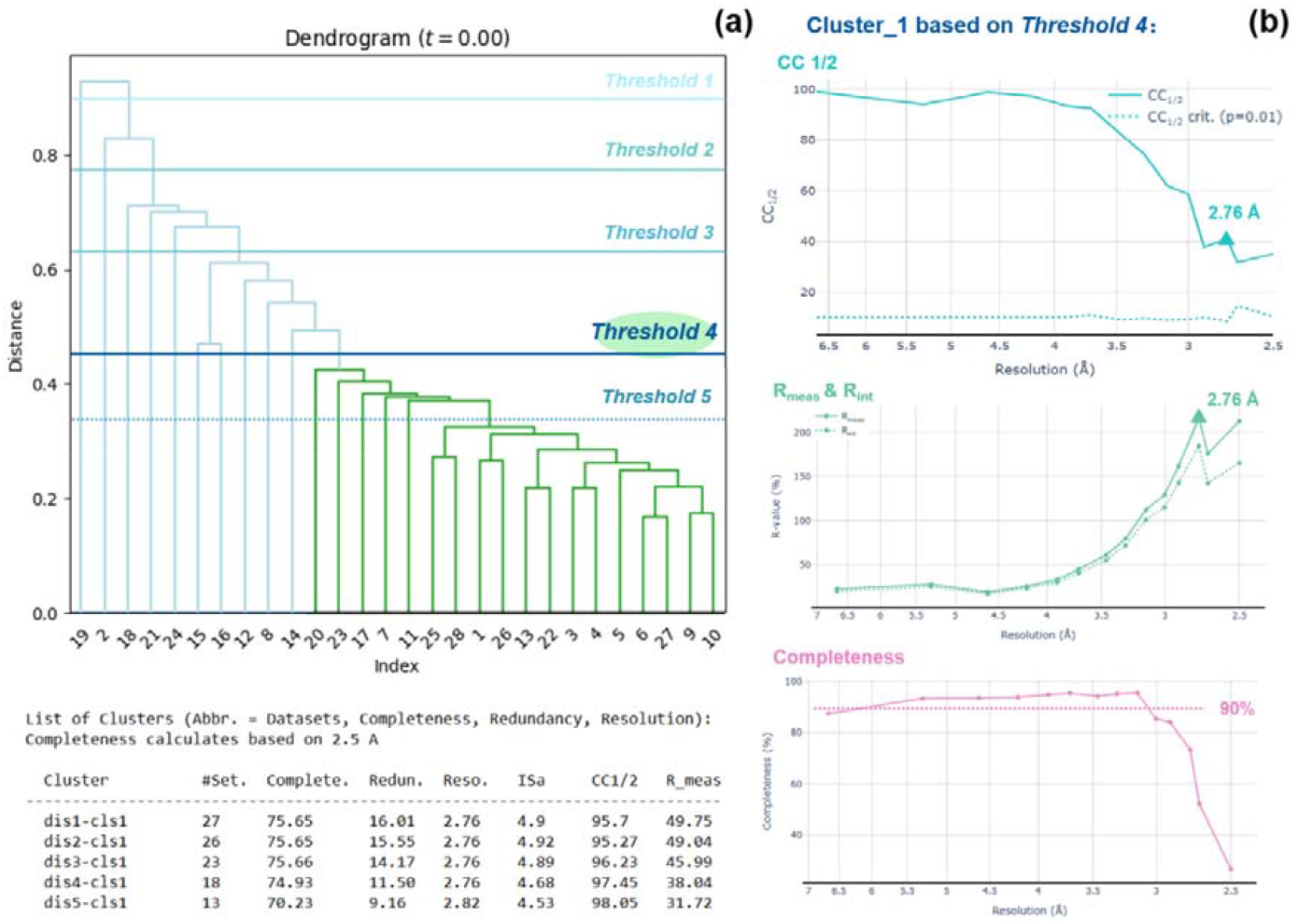
Data processing, analysis and merging of 38 hMTH1 datasets by AutoLEI. (a) Clustering analysis after data merging in the AutoLEI interface. A total of 5 cluster selections were made based on different distances. The summary of the key statistics of these 5 clusters was tabulated in the command window. (b) Web report plots of Cluster_1 based on threshold 4 (18 datasets in total). Plots were captured from the web report.

### 3.4. Real-time data process of triclinic lysozyme

Real-time data processing of single crystal X-ray diffraction is implemented at most major X-ray synchrotron facilities. This capability is also desirable for MicroED, as it could provide live information on current batch data collection, allowing the user to stop collection when enough data has been collected. Moreover, live processing statistics can also provide useful information for optimizing the data collection strategy. By combining AutoLEI with automated data collection protocols, rapid structure determination and phase analysis becomes possible. Here, we used triclinic lysozyme crystals to demonstrate the online MicroED data processing capability of AutoLEI.

Triclinic lysozyme is a challenging sample for MicroED, considering its low symmetry and preferred crystal orientation. Here, the small wedge data collection strategy was also employed to increase data quality. The average tilt range of individual crystals was limited to 15 degrees, making data collection more challenging. As shown in Figure 7a, to achieve real-time data merging (merge mode), unit cell parameters and space group need to be given as prior knowledge. These two input parameters could be easily obtained by first collecting a high-tilt MicroED dataset. In the case of triclinic lysozyme, the expected resolution limit for inspecting live statistics was set to 1.3 Å. Data with *CC*_1/2_ higher than 90% as well as ISa higher than 5 were considered to be good data and used for real-time merging iterations. These parameters could also be customized by editing the “strategy file settings”. Furthermore, the resolution of each good dataset as well as the statistics of merged data completeness and *CC*_1/2_ was updated live. The user can then use this information to check if the current data collection strategy or data filter strategy should be modified.

**Figure 7.**
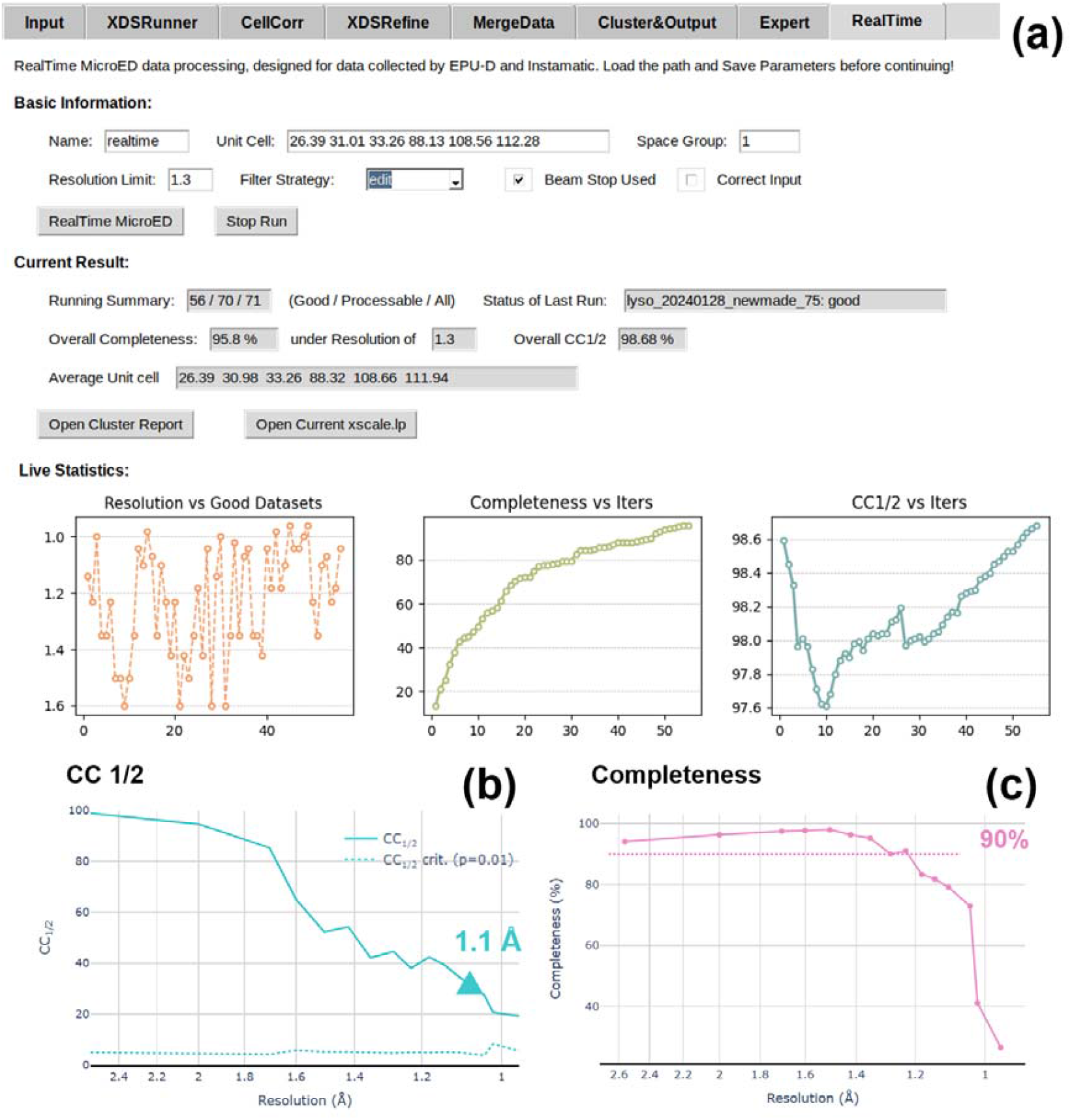
Real-time MicroED data processing of triclinic lysozyme datasets using AutoLEI. (a) The live processing results are displayed under “Live Statistics”. (b) *CC*_1/2_ and (c) Completeness against resolution plots captured from the web report.

After approximately 6 hours of data collection, a total of 71 datasets were collected. The completeness of the data reached 95.8%, while the *CC*_1/2_ was 98.68% at the resolution cut-off of 1.3 Å. Notably, during data collection, the real-time completeness plot revealed a period during which completeness was not significantly improved, indicating preferred crystal orientation. This suggests that the data collection strategy should be altered. We then looked for crystals with different morphologies on the TEM grids. Data collected from those crystals further improved the data completeness. In such an instance, MicroED data collected from high tilting angle might also be considered. The data collection was terminated after the completeness of the merged data reached higher than 95% and the *CC*_1/2_ was above 98% at the pre-set resolution cut-off of 1.3 Å. The web report was then generated to estimate the real high-resolution cut-off. As shown in Figure 7b and 7c, the resolution cut-off was finally set at 1.1 Å, before the *CC*_1/2_ dropped below 30%. The “HKL” file of the final merging iteration was used immediately for phasing and refinement using Phenix (McCoy *et al*., 2007; Afonine *et al*., 2012). The final refinement showed that the *R*_work_ and *R*_free_ converged to 20.42% and 23.45% respectively. The no-filled 2*F*_o_−*F*_c_ electrostatic potential map with the lysozyme structure is presented in Figure 3d.

## 4. Conclusion and outlook

In this paper, we developed an XDS-based GUI, AutoLEI for offline and real-time batch MicroED data processing. We presented four examples utilizing different types of samples to demonstrate the simplicity and usability of AutoLEI, illustrating its potential for fully automated high-throughput structure analyses, such as high-speed ligand screening for structure-based drug discovery and quantitative analysis of heterogenetic phases. The current version of AutoLEI still presents opportunities for further development. We expect to incorporate additional data processing engines, enabling comparative analyses and data integration in subsequent versions. These developments will simplify MicroED data processing and lower the entry barriers for researchers new to the field of MicroED.

## Supporting information

Demonstration - human MTH1

Demonstration - Online data processing

Demonstration - SU-100 MOF

Demonstration - Tyrosine

## Acknowledgements

This work was supported by the European Union’s Horizon 2020 research and innovation programme under the Marie Sklodowska-Curie grant agreement no. 956099 (NanED − Electron Nanocrystallography−H2020-MSCAITN). We also acknowledge financial support from the Swedish Research Council (VR, 2019-00815 (XZ), 2022-03681 (PS) and 2022-03596 (HY)), and the Knut and Alice Wallenberg Foundation (KAW, 2019.0124). We thank Dr. Stef Smith and Dr. Taimin Yang for sharing the code of Edtools. The MicroED data of protein samples were collected at the Cryo-EM Swedish National Facility funded by KAW, Family Erling Persson and Kempe Foundations, SciLifeLab, Stockholm University and Umeå University. We also thank Dr. Mathieu Coinçon, Dr. Dustin Morado and Dr. Marta Carroni for TEM support at Scilifelab.

## Data available

The GUI is available from Zenodo (https://zenodo.org/records/15155988).

The data can be downloaded from Zenodo (https://zenodo.org/records/14536385).

## References

Afonine, P. V., Grosse-Kunstleve, R. W., Echols, N., Headd, J. J., Moriarty, N. W., Mustyakimov, M., Terwilliger, T. C., Urzhumtsev, A., Zwart, P. H. & Adams, P. D. (2012). Acta Crystallogr D Biol Crystallogr 68, 352–367.

Bengtsson, V. E. G., Pacoste, L., De La Rosa-Trevin, J.M., Hofer, G., Zou, X. & Xu, H. (2022). J Appl Crystallogr 55, 638–646.

Brehm, W., Triviño, J., Krahn, J. M., Usón, I. & Diederichs, K. (2023). J Appl Crystallogr 56, 1585–1594.

Cheng, A., Negro, C., Bruhn, J. F., Rice, W. J., Dallakyan, S., Eng, E. T., Waterman, D. G., Potter, C. S. & Carragher, B. (2021). Protein Science 30, 136–150.

Cho, J., Willhammar, T. & Zou, X. (2023). Microporous and Mesoporous Materials 358, 112400.

Cichocka, M. O., Ångström, J., Wang, B., Zou, X. & Smeets, S. (2018). J Appl Crystallogr 51, 1652– 1661.

De La Cruz, M.J., Martynowycz, M. W., Hattne, J. & Gonen, T. (2019). Ultramicroscopy 201, 77–80.

Emsley, P., Lohkamp, B., Scott, W. G. & Cowtan, K. (2010). Acta Crystallogr D Biol Crystallogr 66, 486–501.

Gemmi, M., Mugnaioli, E., Gorelik, T. E., Kolb, U., Palatinus, L., Boullay, P., Hovmöller, S. & Abrahams, J. P. (2019). ACS Cent. Sci. 5, 1315–1329.

Gildea, R. J., Beilsten-Edmands, J., Axford, D., Horrell, S., Aller, P., Sandy, J., Sanchez-Weatherby, J., Owen, C. D., Lukacik, P., Strain-Damerell, C., Owen, R. L., Walsh, M. A. & Winter, G. (2022). Acta Crystallogr D Struct Biol 78, 752–769.

Grape, E. S., Xu, H., Cheung, O., Calmels, M., Zhao, J., Dejoie, C., Proserpio, D. M., Zou, X. & Inge, A. K. (2020). Crystal Growth & Design 20, 320–329.

Guo, P., Shin, J., Greenaway, A. G., Min, J. G., Su, J., Choi, H. J., Liu, L., Cox, P. A., Hong, S. B., Wright, P. A. & Zou, X. (2015). Nature 524, 74–78.

Huang, Z., Grape, E. S., Li, J., Inge, A. K. & Zou, X. (2021). Coordination Chemistry Reviews 427, 213583.

Huang, Z., Willhammar, T. & Zou, X. (2021). Chem. Sci. 12, 1206–1219.

Jemth, A.-S., Gustafsson, R., Bräutigam, L., Henriksson, L., Vallin, K. S. A., Sarno, A., Almlöf, I., Homan, E., Rasti, A., Warpman Berglund, U., Stenmark, P. & Helleday, T. (2018). Nucleic Acids Research 10.1093/nar/gky896.

Kabsch, W. (2010). Acta Crystallogr D Biol Crystallogr 66, 125–132.

Kolb, U., Gorelik, T., Kübel, C., Otten, M. T. & Hubert, D. (2007). Ultramicroscopy 107, 507–513.

Liebschner, D., Afonine, P. V., Baker, M. L., Bunkóczi, G., Chen, V. B., Croll, T. I., Hintze, B., Hung, L.-W., Jain, S., McCoy, A. J., Moriarty, N. W., Oeffner, R. D., Poon, B. K., Prisant, M. G., Read, R. J., Richardson, J. S., Richardson, D. C., Sammito, M. D., Sobolev, O. V., Stockwell, D. H., Terwilliger, T. C., Urzhumtsev, A. G., Videau, L. L., Williams, C. J. & Adams, P. D. (2019). Acta Crystallogr D Struct Biol 75, 861–877.

Lightowler, M., Li, S., Ou, X., Cho, J., Liu, B., Li, A., Hofer, G., Xu, J., Yang, T., Zou, X., Lu, M. & Xu, H. (2024). Angewandte Chemie International Edition 63, e202317695.

Lightowler, M., Li, S., Ou, X., Zou, X., Lu, M. & Xu, H. (2022). Angew Chem Int Ed 61, e202114985.

Lu, P., Xu, J., Sun, Y., Guillet-Nicolas, R., Willhammar, T., Fahda, M., Dib, E., Wang, B., Qin, Z., Xu, H., Cho, J., Liu, Z., Yu, H., Yang, X., Lang, Q., Mintova, S., Zou, X. & Valtchev, V. (2024). Nature 636, 368– 373.

Luo, Y., Wang, B., Smeets, S., Sun, J., Yang, W. & Zou, X. (2023). Nat. Chem. 10.1038/s41557-022-01131-8.

Martynowycz, M. W., Clabbers, M. T. B., Hattne, J. & Gonen, T. (2022). Nat Methods 19, 724–729.

McCoy, A. J., Grosse-Kunstleve, R. W., Adams, P. D., Winn, M. D., Storoni, L. C. & Read, R. J. (2007). J Appl Crystallogr 40, 658–674.

Palatinus, L., Brázda, P., Jelínek, M., Hrdá, J., Steciuk, G. & Klementová, M. (2019). Acta Crystallogr B Struct Sci Cryst Eng Mater 75, 512–522.

Samperisi, L., Jaworski, A., Kaur, G., Lillerud, K. P., Zou, X. & Huang, Z. (2021). J. Am. Chem. Soc. 143, 17947–17952.

Shi, D., Nannenga, B. L., Iadanza, M. G. & Gonen, T. (2013). eLife 2, e01345.

Truong, K.-N., Ito, S., Wojciechowski, J. M., Göb, C. R., Schürmann, C. J., Yamano, A., Del Campo, M., Okunishi, E., Aoyama, Y., Mihira, T., Hosogi, N., Benet-Buchholz, J., Escudero-Adán, E. C., White, F. J., Ferrara, J. D. & Bücker, R. (2023). Symmetry 15, 1555.

Unge, J., Lin, J., Weaver, S. J., Sae Her, A. & Gonen, T. (2024). Advanced Science 11, 2400081.

Van Genderen, E., Clabbers, M. T. B., Das, P. P., Stewart, A., Nederlof, I., Barentsen, K. C., Portillo, Q., Pannu, N. S., Nicolopoulos, S., Gruene, T. & Abrahams, J. P. (2016). Acta Crystallogr A Found Adv 72, 236–242.

Wan, W., Sun, J., Su, J., Hovmöller, S. & Zou, X. (2013). J Appl Crystallogr 46, 1863–1873.

Wang, B., Zou, X. & Smeets, S. (2019). IUCrJ 6, 854–867.

Winter, G., Beilsten-Edmands, J., Devenish, N., Gerstel, M., Gildea, R. J., McDonagh, D., Pascal, E., Waterman, D. G., Williams, B. H. & Evans, G. (2022). Protein Science 31, 232–250.

Winter, G., Waterman, D. G., Parkhurst, J. M., Brewster, A. S., Gildea, R. J., Gerstel, M., Fuentes-Montero, L., Vollmar, M., Michels-Clark, T., Young, I. D., Sauter, N. K. & Evans, G. (2018). Acta Crystallogr D Struct Biol 74, 85–97.

Xu, H., Lebrette, H., Clabbers, M. T. B., Zhao, J., Griese, J. J., Zou, X. & Högbom, M. (2019). Sci. Adv. 5, eaax4621.

Xu, H., Lebrette, H., Yang, T., Srinivas, V., Hovmöller, S., Högbom, M. & Zou, X. (2018). Structure 26, 667-675.e3.

Yoshimura, D., Sakumi, K., Ohno, M., Sakai, Y., Furuichi, M., Iwai, S. & Nakabeppu, Y. (2003). Journal of Biological Chemistry 278, 37965–37973.

Zhang, D., Oleynikov, P., Hovmöller, S. & Zou, X. (2010). Zeitschrift Für Kristallographie 225, 94–102.

