## Supplementary material for "AutoLEI: An XDS-based Graphical User Interface for Automated Real-time and Offline Batch 3D ED/MicroED Data Processing": SI document

### Supporting information

#### S1. Supporting function implementation in AutoLEI

##### S1.1 *XDSRunner*

*Format Transfer:* The format converter leverages functions in Python libraries: mrcfile (Burnley *et al.*, 2017), FabIO (Knudsen *et al.*, 2013), and h5py (https://www.h5py.org/). When converting MRC files to IMG files, a pedestal is added to prevent negative pixel values. The value of the pedestal was dynamically global and the calculation was based on the lowest average (negative) intensity of a random 10 frames in the datasets.

*Find Beam Centre:* This function identifies the most intense region in the image and designates its center as the position of the direct beam.

*Find Beam Stop:* Using the diffuse scattering of the direct beam, this function determines the beam center and identifies the shadow of the beam stop. It designates the beam stop location as an untrusted area in XDS processing. Image processing for this task employs functions in Skimage (Van Der Walt *et al.*, 2014).

*Estimate resolution:* The resolution of a single/merged dataset is determined by $CC_{1/2}$, $R_{int}$ and I/Sigma. Practically, the $CC_{1/2}$ should be statistically significant, and the $R_{int}$ should be less than 180%. When the $R_{int}$ is 100–180%, the I/Sigma of the corresponding shell should be above 0.5.

*Estimate Symmetry & Cell-Cluster:*

*Estimate Symmetry:* The lattice symmetry, determined solely from cell parameters, represents the fundamental symmetry of the structure and is always equal to or higher than the actual symmetry derived from intensities. AutoLEI processes the CORRECT.LP output from XDS to extract possible lattice symmetries. It then utilizes its internal HKL analyzer to calculate $R_{\text{meas}}$ and $CC_{1/2}$ values for potential Laue groups. Four indicators are provided for each Laue group: Figure of merit (FOM) from XDS, Lattice deviation from AutoLEI, $R_{\text{meas}}$ ratio and $CC_{1/2}$ ratio. The definitions of $R_{\text{meas}}$ ratio and $CC_{1/2}$ ratio are:

$$R_{\text{meas}}\ \text{ratio} = R_{\text{meas}}(\text{Laue})/R_{\text{meas}}(P1)$$

$$CC_{1/2}\ \text{ratio} = [1 - CC_{1/2}(\text{Laue})]/[1 - CC_{1/2}(P1)]$$

The $CC_{1/2}$ ratio is designed to address intensity inconsistencies in symmetry-related reflections caused by random and systematic errors, including the inevitable dynamical effects of electron diffraction, which is more significant in crystals of higher symmetry.

*Cell-Cluster*: The initial choice of a crystallographic unit cell is arbitrary. Therefore, Niggli cell reduction is employed to transform cell parameters into a unique, standardized form defined by rigorous mathematical conditions. This ensures a one-to-one correspondence between the lattice and its reduced-cell representation (Niggli, 1928). According to Buerger's cell (also known as the $G^6$-tensor) (Buerger, 1957), Niggli cells are categorized into 44 types (De Wolff & Gruber, 1991). The two main types, Type I and Type II, are determined by the sign of the diagonal elements of the G-tensor. Type-I have angles less than 90° while Type-II have angles greater than 90°. In triclinic (*aP*), monoclinic (*mC*), and hexagonal (*hP*) systems, there are simultaneously two valid settings of the unit cell corresponding to Type I and Type II.

In borderline cases, similar lattices may correspond to two different types of Niggli-reduced cells within experimental error. For example, *aP* lattices (10.1, 14.2, 15.3, 91.1, 108.7, 110.7) and (10.1, 14.21, 15.26, 75.6, 70.7, 69.2) are both primary Niggli reduced cell for *mI* lattice (18.1, 10.1, 21.0, 90, 95.04, 90). *cF* lattice (4.48, 4.48, 4.48, 90, 90, 90) can also have different Niggli reduced cells, such as (3.162, 3.163, 3.164, 60.11, 89.84, 60.05) and (3.162, 3.163, 3.164, 90.19, 119.74, 119.82). Those instances indicate the presence of latent higher symmetry, as the lattice parameters are close to those required for a higher symmetry space group or a more restrictive Bravais lattice type (Grosse-Kunstleve, 1999). This ambiguity can cause some unit-cell clustering methods to fail, as they rely solely on metric comparisons of primary unit cells. Therefore, recognizing and properly classifying these subtle cases is essential for accurate symmetry determination. In this program, we address these challenges by performing unit-cell clustering on cells with "prompted" symmetry, ensuring that even borderline configurations are correctly identified and correlated with their underlying crystallographic symmetries.

After estimating the symmetry, the lattice symmetry is extracted from the data reduction results of XDS. Before calculating the distance between unit cells, a Niggli reduction is first performed based on previously described methodology (Grosse-Kunstleve *et al.*, 2004). To accurately reflect the measurement errors inherent in the electron diffraction pattern, the unit cell is transformed into a matrix in reciprocal space, where Procrustes Analysis is applied to calculate the distance between lattices. To prevent over scaling of the reciprocal cell matrix, the distance between unit cells is adjusted by the ratio of their volumes in an exponential format. The final distance is defined as:

$$D = d(\text{Procrustes}, M_1, M_2) \times \exp\left(\frac{V_1}{V_2}\right), \text{where } V = \frac{1}{\det M}$$

The resulting distance matrix is then used for cluster analysis. If specified in the settings file, clusters are further refined based on the highest symmetry. For reporting purposes, within each cluster, all possible lattice symmetries are summarized, and their metrics are averaged.

##### S1.2 *XDSRefine*

*Rotation Axis*: The algorithm is adopted with enhanced efficiency and accuracy from *edtools* (https://github.com/instamatic-dev/edtools), using its original code from a previous publication (Kolb *et al.*, 2012).

*Divergence, Mosaicity and Scale*: These parameters are extracted from INTEGRATE.LP. The average value is calculated by omitting outliers. It may be different from the recommended value from XDS.

*Beam Centre*: An algorithm is developed to load reflection points from SPOT.XDS and then estimate the beam center by constructing as many Friedel pairs as possible.

*Index Ratio*: This function will run IDXREF with different parameters and use the best combination of input parameters to continue.

*View Reciprocal Space*: This function would read SPOT.XDS from XDS and reconstruct the reciprocal space without considering Ewald correction.

##### S1.3 *MergeData*

*Merge data*: AutoLEI will only merge data in separate datasets within the resolution limit. In the AutoLEI workflow, the resolution is not cut during processing, instead, the resolution is cut when merging. This process avoids weak/noisy/wrong reflections merged into the final "HKL" file.

##### S1.4 *Cluster&Output*

*Intensity-Cluster*: The distance is calculated from the correlation coefficient between datasets in XSCALE.LP from XSCALE. The algorithm was described in a previous publication (Giordano *et al.*, 2012).

$$d = \sqrt{1 - CC^2}$$

#### S2. Experiments

##### Sample preparation for tyrosine and SU-100

Commercial L-Tyrosine powder (Sigma-Aldrich) was ground using two glass plates. The crushed powder was then loaded onto a R2/2 Cu300 Quantifoil TEM grid.

SU-100 was synthesized as previously described (Grape *et al.*, 2020). The as-synthesized SU-100 powder was first crushed using a mortar and then dispersed in water. The R2/2 Cu300 Quantifoil TEM grid was rapidly dipped into a 1% Tween-20 solution to make the TEM grid hydrophilic. Filter paper was used to remove the extra Tween-20 solution by back blotting. The SU-100 suspension was then transferred onto the treated TEM grid.

##### Sample preparation for hMTH1 and lysozyme proteins

The expression and purification of protein hMTH1 was performed as described previously (Svensson *et al.*, 2011). The precipitant solution used for hMTH1 crystallization contained 30% PEG6000, 0.1 M sodium acetate buffer (pH 4) and 0.16 M lithium sulfate. After mixing 0.5 μL protein solution (14 mg/mL) and 0.5 μL precipitant solution, needle-like crystals formed within 30 mins. A 1 μL volume of these crystals was then mixed with 15 μL of precipitant solution and ground into small fragments. 15 μL of protein solution was then added into the mixture to produce the micro-sized crystals used in this study.

Lyophilized lysozyme powder (Sigma-Aldrich, 62970) from chicken egg white was dissolved in 50 mM sodium acetate buffer (pH 4.5) to make a 20 mg/mL protein solution. A precipitant solution containing 0.4 M sodium nitrate and 50 mM sodium acetate (pH 4.5) was also prepared. For crystallization, 30 μL of both protein solution and precipitant solution were added into a PCR tube. The tube was sealed with parafilm and stored overnight at 4 °C (Wang *et al.*, 2007). The tube was then stored at room temperature. After 3-4 days, several X-ray-sized crystals could be observed in the PCR tube. To obtain crystals used for MicroED data collection, these crystals were ground and used as seeds. 1 μL of seed crystal slurry was added to 20 μL of precipitant solution. Microcrystals were formed by mixing the slurry with protein solution.

The grids for both hMTH1 and lysozyme were prepared as follows: the R1.2/1.3 Cu300 Quantifoil grid was firstly dipped into a 1% Tween-20 solution and back-blotted using filter paper to make the carbon supporting film hydrophilic. A 1.5 μL drop of crystal slurry was then pipetted onto the treated TEM grid. The grid was then back-blotted again to remove excess liquid and plunge frozen in liquid ethane. The grids were then clipped and transferred into the autoloader of a Titan Krios cryo-TEM (ThermoFisher Scientific) under cryogenic conditions.

##### Data collection

MicroED data of tyrosine and SU-100 were collected using a JEOL JEM-2100 transmission electron microscope equipped with a Timepix detector (Amsterdam Scientific Instrument), operated at 200 keV.

A Gatan 914 cryo-transfer holder was used to collect data at cryogenic temperature (100 K). *Instamatic* (Cichocka *et al.*, 2018) was used for high-throughput data collection with crystal tracking and metadata recording. The tilt step for data collection was 0.23 degrees/frame and the exposure time was 0.5 s/frame. The tilting angle could be observed in the AutoLEI report.

MicroED data collection of hMTH1 protein micro-crystals was performed using a Titan Krios G3i microscope (300 keV) while lysozyme data was collected on a Titan Krios G2 microscope (300 keV). Both microscopes were equipped with an autoloader for sample loading and a Ceta-D detector for data collection. EPUD was used for data collection. In both cases, data were collected under nanoprobe mode, without the use of a selected area aperture. The beam size used for hMTH1 and lysozyme data collection was 1 μm and 1.5 μm, while the flux used for data collection was 0.14 $e^-/Å^2/s$ and 0.12 $e^-/Å^2/s$, respectively. The oscillation range used for hMTH1 data collection was 1°/frame and the exposure time was 1 s/frame. The total tilting range for each dataset was 15 or 20 degrees. For lysozyme, the oscillation range was set to 0.5°/frame and the exposure time was set to 1 s/frame. The total tilt range for each dataset was 15 degrees.

#### Structure determination

Structure determination of L-tyrosine and SU-100 was performed in Olex2 (Dolomanov *et al.*, 2009), using SHELXS for structure solution and SHELXL (Sheldrick, 2008) for structure refinement. The structures of hMTH1 and lysozyme proteins were determined using Phenix (Liebschner *et al.*, 2019). Molecular replacement was carried out using Phaser (McCoy *et al.*, 2007) and the structure refinement was performed with Phenix.refine (Afonine *et al.*, 2012).

##### S3. AutoLEI graphical user interface

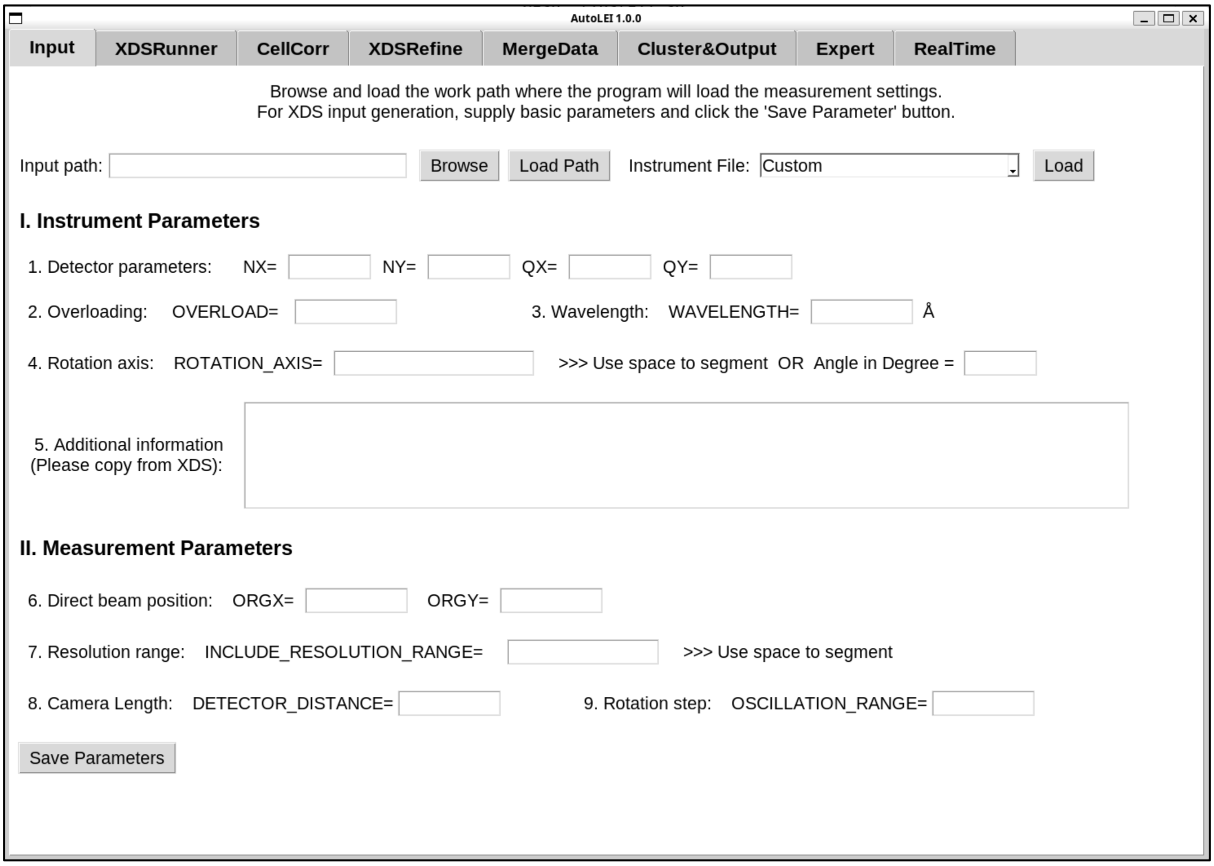


**Figure S1** AutoLEI interface of ***Input***

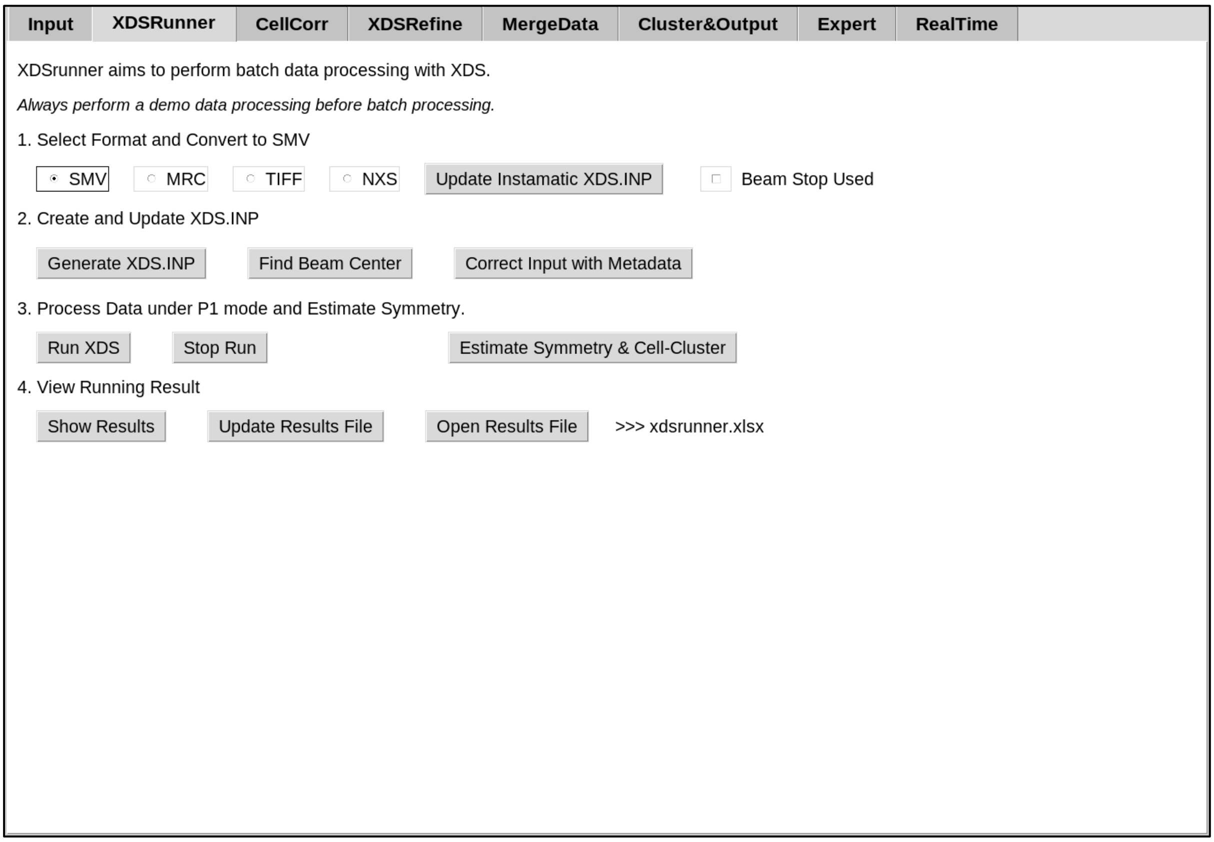


**Figure S2** AutoLEI interface of ***XDSRunner***

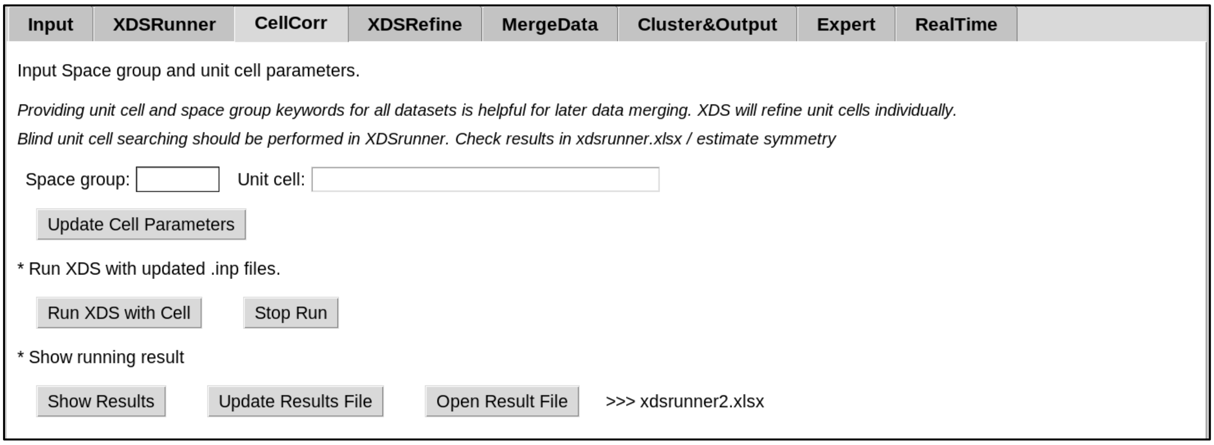


**Figure S3** AutoLEI interface of ***CellCorr***

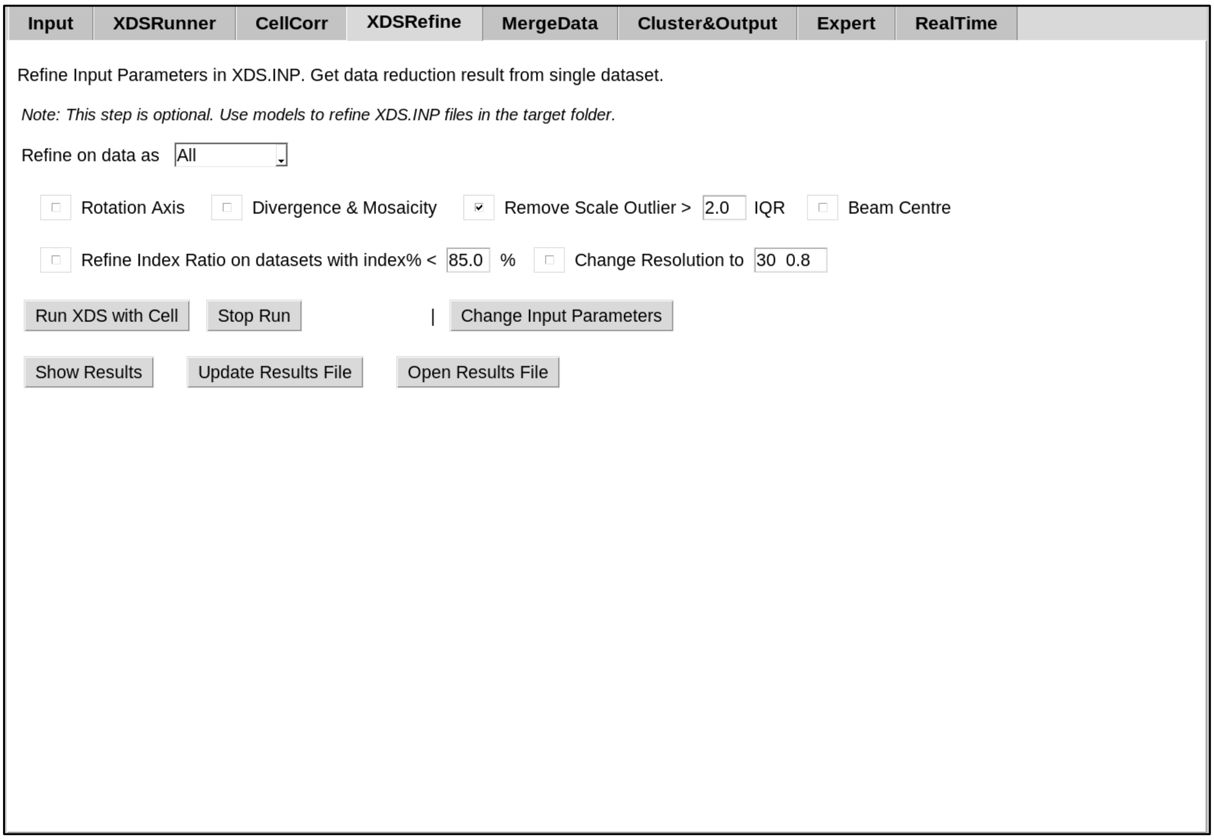


**Figure S4** AutoLEI interface of ***XDSRefine***

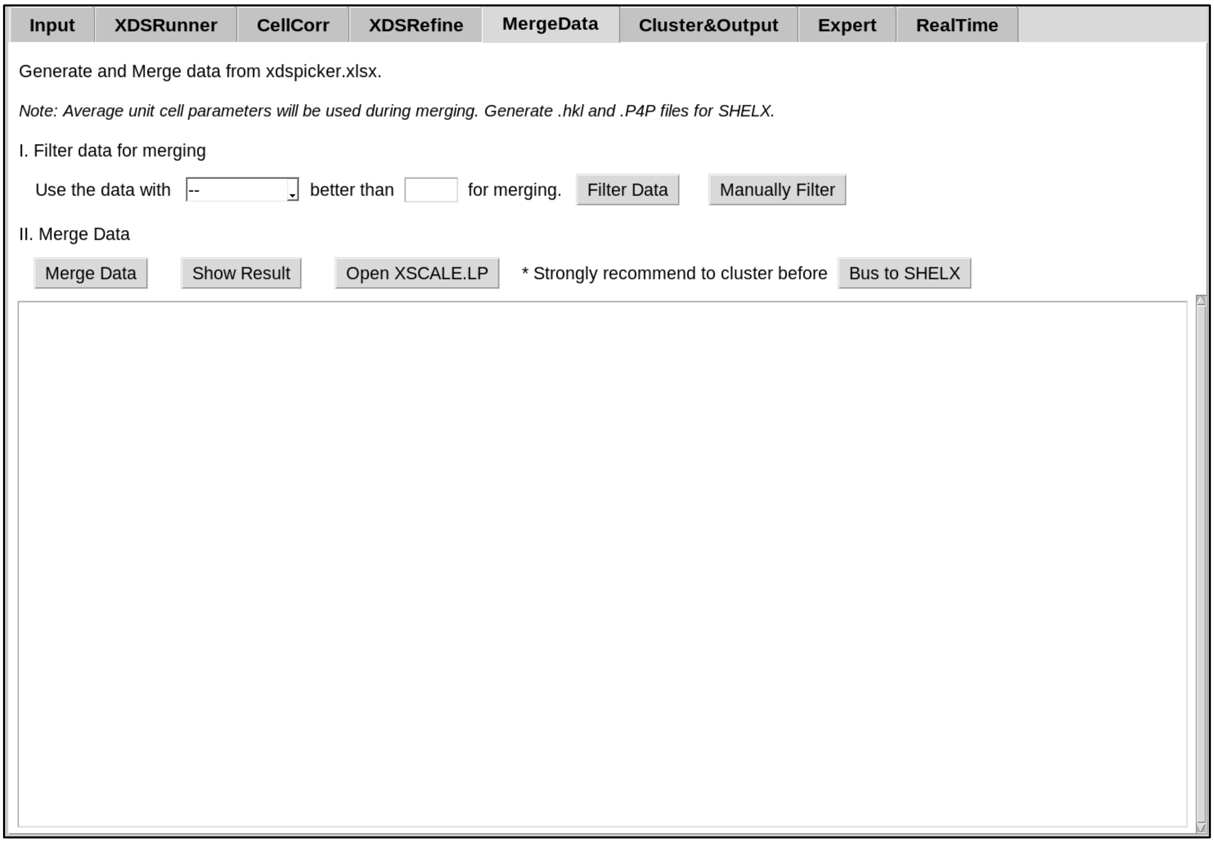


**Figure S5** AutoLEI interface of ***MergeData***

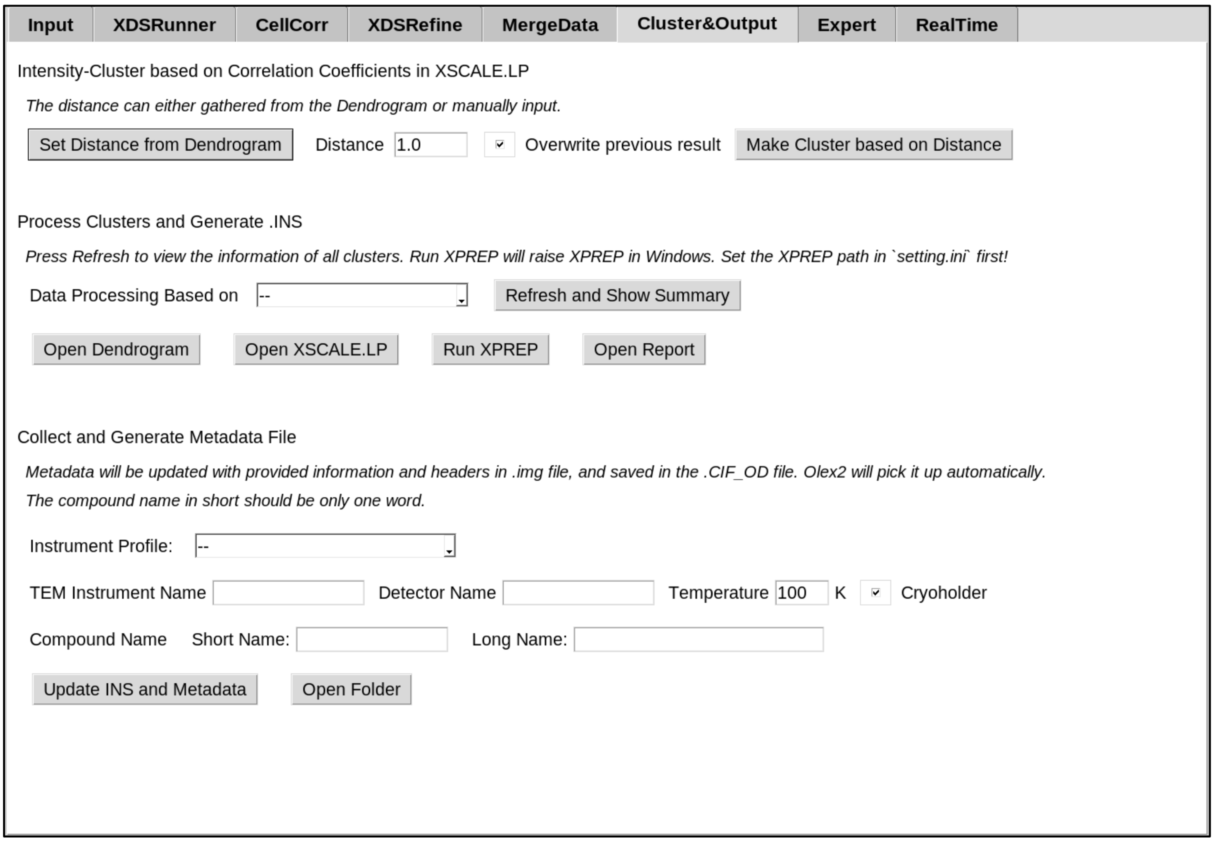


**Figure S6** AutoLEI interface of ***Cluster***

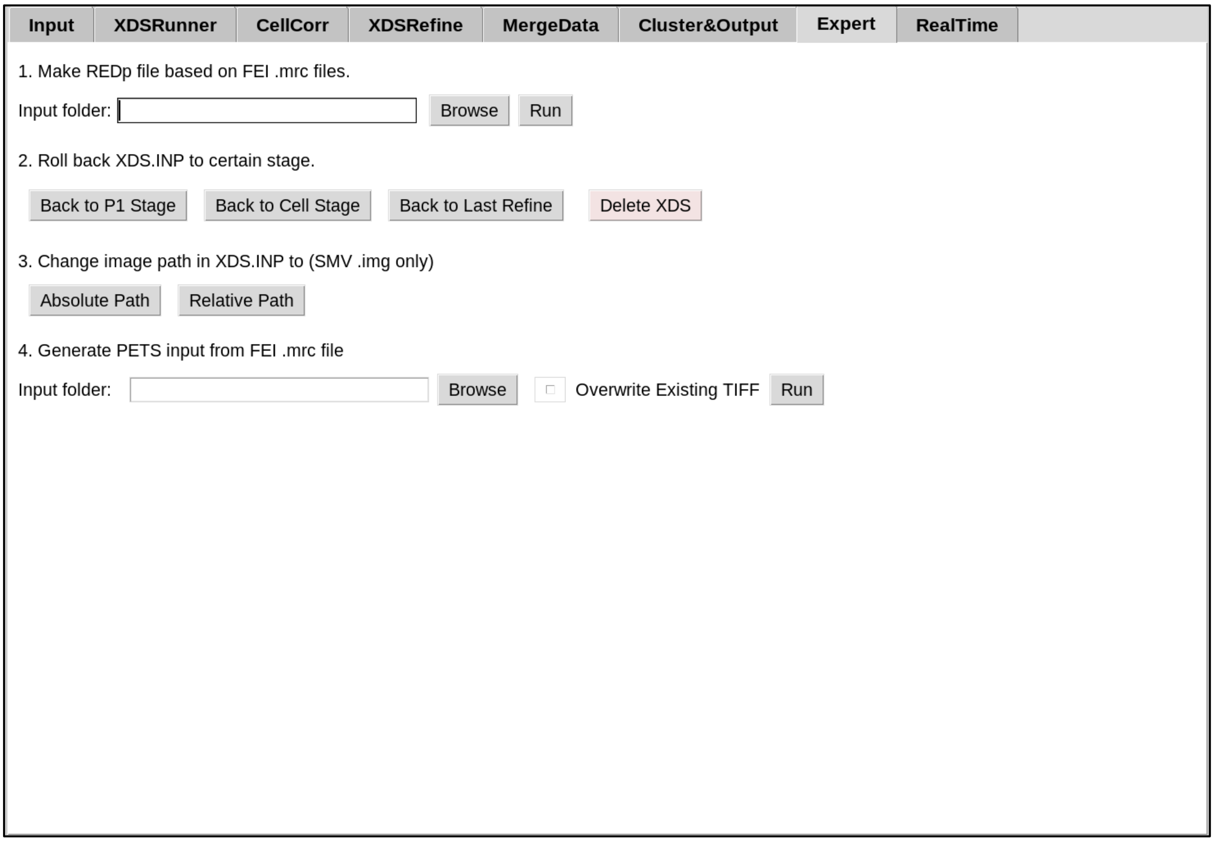


**Figure S7** AutoLEI interface of ***Expert***

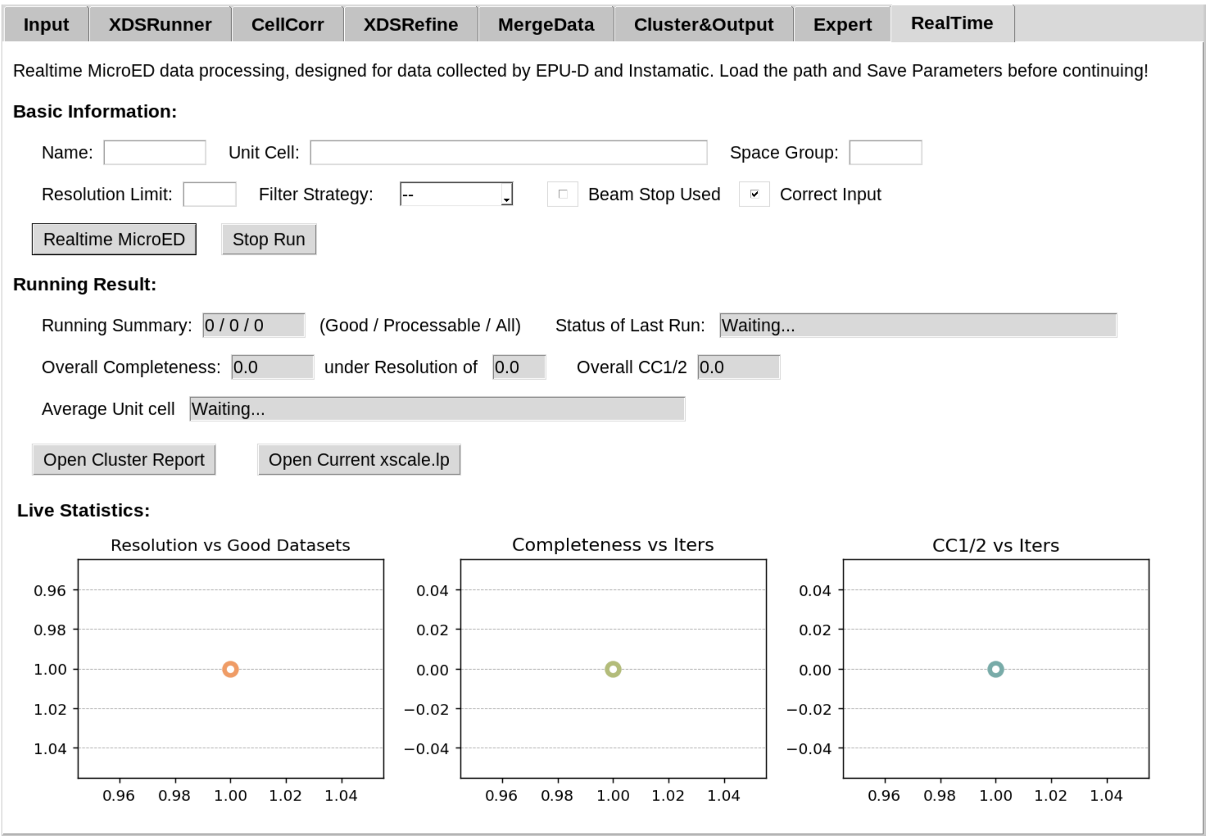


**Figure S8** AutoLEI interface of ***RealTime***

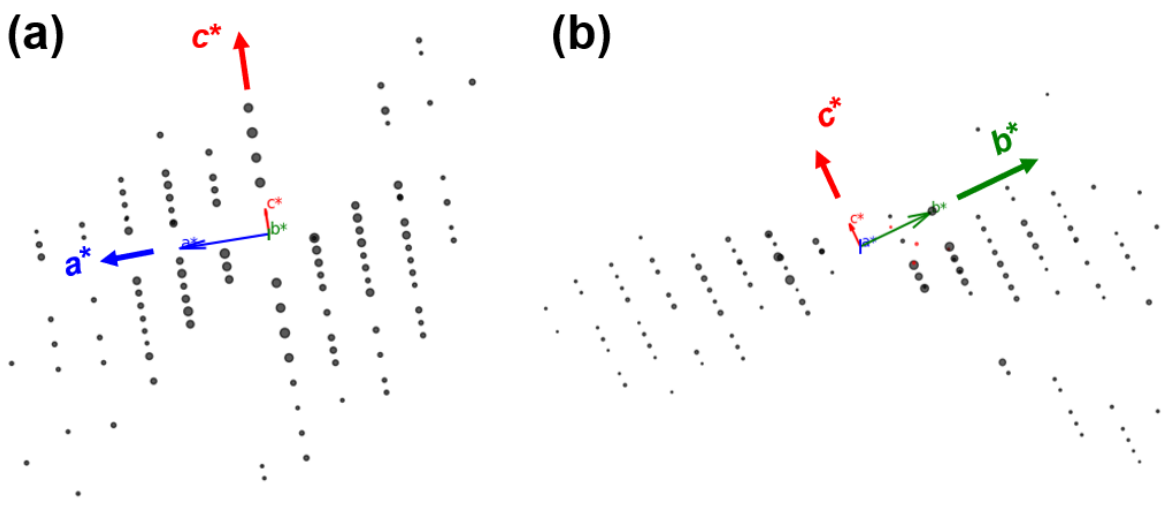


**Figure S9** Systematic absence of L-tyrosine in reciprocal space (a) *h*0*l* slice of 3D reciprocal data from Experiment_10 and (b) 0*kl* slice of 3D reciprocal data from Experiment_12. The reflection conditions could be deduced as *h*00: *h* = 2n, 0*k*0: *k* = 2n and 00*l*: *l* = 2n.

#### S4. Crystallographic tables of the four different samples

**Table S1** Crystallographic data and refined parameters data of 12 tyrosine datasets

| No. | Resolution (Å) | Completeness (%) | ISa | $R_{meas}$ (%) | CC1/2 (%) | Space group | *a* (Å) | *b* (Å) | *c* (Å) | No. of unique reflections | No. of total reflections | $R_{int}$ | No. of parameters | No. of restraints | $R_1$ (obs) | $R_1$ (all) | $wR_2$ (all) | GOOF |
|---|---|---|---|---|---|---|---|---|---|---|---|---|---|---|---|---|---|---|
| 1 | 0.56 | 60.01 | 9.33 | 15.38 | 99.10 | $P2_12_12_1$ | 5.994(8) | 7.076(4) | 21.745(4) | 2933 | 5214 | 0.1140 | 162 | 8 | 0.1578 | 0.2059 | 0.4246 | 1.220 |
| 2 | 0.55 | 57.16 | 11.5 | 12.79 | 99.10 | $P2_12_12_1$ | 5.997(6) | 7.035(5) | 21.796(9) | 2867 | 5333 | 0.0940 | 151 | 14 | 0.1587 | 0.2076 | 0.4041 | 1.270 |
| 3 | | | | | | | | | No solution | | | | | | | | | |
| 4 | 0.56 | 50.29 | 7.45 | 17.07 | 98.70 | $P2_12_12_1$ | 5.997(12) | 6.984(3) | 21.761(7) | 2453 | 4932 | 0.1331 | 162 | 10 | 0.1398 | 0.1955 | 0.3756 | 1.048 |
| 5* | 0.74 | 74.37 | 5.09 | 24.64 | 96.37 | $P2_12_12_1$ | 6.011(16) | 7.029(6) | 21.75(6) | 2477 | 5119 | 0.1900 | 54 | 0 | 0.2036 | 0.2766 | 0.4854 | 1.198 |
| 6 | 0.56 | 49.63 | 6.33 | 22.16 | 97.30 | $P2_12_12_1$ | 5.998(19) | 6.981(14) | 21.732(17) | 2324 | 3530 | 0.1843 | 158 | 21 | 0.1504 | 0.2125 | 0.4027 | 1.003 |
| 7 | | | | | | | | | No solution | | | | | | | | | |
| 8 | 0.54 | 58.21 | 6.42 | 18.45 | 97.80 | $P2_12_12_1$ | 6.017(16) | 7.049(5) | 21.752(14) | 3204 | 5423 | 0.1399 | 158 | 34 | 0.1605 | 0.2059 | 0.4001 | 1.093 |
| 9* | 0.54 | 53.01 | 8.31 | 14.61 | 99.10 | $P2_12_12_1$ | 5.995(9) | 6.957(13) | 21.827(16) | 2694 | 4903 | 0.1089 | 55 | 0 | 0.2161 | 0.3065 | 0.5664 | 1.666 |
| 10 | 0.68 | 87.61 | 7.51 | 19.45 | 98.44 | $P2_12_12_1$ | 6.0080(12) | 7.0250(14) | 21.690(4) | 2592 | 5506 | 0.1558 | 162 | 8 | 0.1156 | 0.1764 | 0.3097 | 0.998 |
| 11 | 0.56 | 45.46 | 11.15 | 11.18 | 98.8 | $P2_12_12_1$ | 5.988(4) | 7.193(13) | 21.733(5) | 2100 | 3868 | 0.0881 | 143 | 10 | 0.1596 | 0.1857 | 0.4111 | 1.419 |
| 12 | 0.56 | 85.16 | 7.19 | 24.21 | 97.60 | $P2_12_12_1$ | 6.010(17) | 7.050(7) | 21.77(5) | 3199 | 6209 | 0.1892 | 162 | 13 | 0.1505 | 0.1955 | 0.3917 | 1.083 |

[*] Isotropic refinement for non-hydrogen atoms.

**Table S2** Crystallographic data and refined parameters data of SU-100

| **SU-100** | |
|---|---|
| Chemical Formula | $C_{15}H_7BiO_6$ |
| No. datasets | 11 |
| Wavelength (Å) | 0.02508 |
| Temperature (K) | 100 |
| Space group | *C*2/*c* |
| *a* (Å) | 26.44(3) |
| *b* (Å) | 10.049(17) |
| *c* (Å) | 18.135(17) |
| *β* (°) | 127.28(9) |
| Volume (Å$^3$) | 3834(9) |
| Z | 8 |
| Completeness (%, to 0.84 Å) | 94.9 |
| Resolution (Å) | 0.62 |
| $R_{meas}$ (%) | 44.3 |
| CC1/2 (%) | 97.8 |
| $R_{int}$ | 0.3566 |
| No. unique reflections | 7193 |
| No. total reflections | 95728 |
| No. parameters | 200 |
| No. restraints | 0 |
| R1 (obs) | 0.1946 |
| R1 (all) | 0.2572 |
| wR2 (all) | 0.4865 |
| GOOF | 1.103 |

**Table S3** Crystallographic data and refined parameters data of hMT1 and lysozyme

| **Sample** | **apo-hMTH1** | **Lysozyme** |
|---|---|---|
| **Data processing** | | |
| No. of crystals | 18 | 56 |
| Space group | $P2_12_12_1$ | $P1$ |
| $a$, $b$, $c$ (Å) | 61.66, 69.71, 80.87 | 26.39, 30.98, 33.26 |
| $\alpha$, $\beta$, $\gamma$ (°) | 90, 90, 90 | 88.298, 108.613, 111.985 |
| Resolution (Å) | 23.13 – 2.76 | 10.53 – 1.1 |
| CC1/2 (%) | 97.9 (42.8) | 98.6 (31.5) |
| $R_{merge}$ (%) | 44.2 (151.8) | 30.8 (118.9) |
| $R_{meas}$ (%) | 45.9 (164.0) | 33.1 (134.1) |
| $R_{pim}$ (%) | 11.6 (56.2) | 11.3 (58.3) |
| Multiplicity | 12.5 (6.5) | 7.0 (4.6) |
| Completeness (%) | 90.9 (75.2) | 91.1 (78.7) |
| **Refinement** | | |
| No. unique reflections | 8397 | 34030 |
| $R_{work}/R_{free}$ | 21.07/26.82 | 20.42/23.45 |
| Mean B-factors (Å$^2$) | 39.01 | 9.68 |
| R.M.S. deviations | | |
| Bond lengths (Å) | 0.002 | 0.023 |
| Bond angle (°) | 0.549 | 1.484 |
| Ramachandran | | |
| Favoured (%) | 95.08 | 98.40 |
| Allowed (%) | 4.59 | 1.6 |
| Outliers (%) | 0.33 | 0 |
| Clashscore | 2.01 | 7.21 |
| Rotamer outliers (%) | 1.11 | 0 |

Values in parentheses are for the highest-resolution shell

#### Reference


Afonine, P. V., Grosse-Kunstleve, R. W., Echols, N., Headd, J. J., Moriarty, N. W., Mustyakimov, M., Terwilliger, T. C., Urzhumtsev, A., Zwart, P. H. & Adams, P. D. (2012). *Acta Crystallogr D Biol Crystallogr* **68**, 352–367.

Buerger, M. J. (1957). Z. Kristallogr. Cryst. Mater. 109, 42–60.

Burnley, T., Palmer, C. M. & Winn, M. (2017). *Acta Crystallogr D Struct Biol* **73**, 469–477.

Cichocka, M. O., Ångström, J., Wang, B., Zou, X. & Smeets, S. (2018). *J Appl Crystallogr* **51**, 1652–1661.

De Wolff, P. M. & Gruber, B. (1991). *Acta Crystallogr A Found Crystallogr* **47**, 29–36.

Dolomanov, O. V., Bourhis, L. J., Gildea, R. J., Howard, J. A. K. & Puschmann, H. (2009). *J Appl Crystallogr* **42**, 339–341.

Giordano, R., Leal, R. M. F., Bourenkov, G. P., McSweeney, S. & Popov, A. N. (2012). *Acta Crystallogr D Biol Crystallogr* **68**, 649–658.

Grape, E. S., Xu, H., Cheung, O., Calmels, M., Zhao, J., Dejoie, C., Proserpio, D. M., Zou, X. & Inge, A. K. (2020). *Crystal Growth & Design* **20**, 320–329.

Grosse-Kunstleve, R. W. (1999). *Acta Crystallogr A Found Crystallogr* **55**, 383–395.

Grosse-Kunstleve, R. W., Sauter, N. K. & Adams, P. D. (2004). *Acta Crystallogr A Found Crystallogr* **60**, 1–6.

Knudsen, E. B., Sørensen, H. O., Wright, J. P., Goret, G. & Kieffer, J. (2013). *J Appl Crystallogr* **46**, 537–539.

Kolb, U., Shankland, K., Meshi, L., Avilov, A. & David, W. I. F. (2012). Uniting Electron Crystallography and Powder Diffraction Dordrecht: Springer Netherlands.

Liebschner, D., Afonine, P. V., Baker, M. L., Bunkóczi, G., Chen, V. B., Croll, T. I., Hintze, B., Hung, L.-W., Jain, S., McCoy, A. J., Moriarty, N. W., Oeffner, R. D., Poon, B. K., Prisant, M. G., Read, R. J., Richardson, J. S., Richardson, D. C., Sammito, M. D., Sobolev, O. V., Stockwell, D. H., Terwilliger, T. C., Urzhumtsev, A. G., Videau, L. L., Williams, C. J. & Adams, P. D. (2019). *Acta Crystallogr D Struct Biol* **75**, 861–877.

Niggli, P. 1928. Handbuch der Experimentalphysik, Vol. 7, Part 1, pp. 108-176. Leipzig: Akademische Verlagsgesellschaft

McCoy, A. J., Grosse-Kunstleve, R. W., Adams, P. D., Winn, M. D., Storoni, L. C. & Read, R. J. (2007). *J Appl Crystallogr* **40**, 658–674.

Sheldrick, G. M. (2008). *Acta Crystallogr A Found Crystallogr* **64**, 112–122.

Svensson, L. M., Jemth, A.-S., Desroses, M., Loseva, O., Helleday, T., Högbom, M. & Stenmark, P. (2011). *FEBS Letters* **585**, 2617–2621.

Van Der Walt, S., Schönberger, J. L., Nunez-Iglesias, J., Boulogne, F., Warner, J. D., Yager, N., Gouillart, E. & Yu, T. (2014). *PeerJ* **2**, e453.

Wang, J., Dauter, M., Alkire, R., Joachimiak, A. & Dauter, Z. (2007). *Acta Crystallogr D Biol Crystallogr* **63**, 1254–1268.
